# Acidity and sulfur oxidation intermediate concentrations controlled by O_2_-driven partitioning of sulfur oxidizing bacteria in a mine tailings impoundment

**DOI:** 10.1101/2021.09.16.460096

**Authors:** Kelly J. Whaley-Martin, Lin-Xing Chen, Tara Colenbrander Nelson, Jennifer Gordon, Rose Kantor, Lauren E. Twible, Stephanie Marshall, Laura Rossi, Benoit Bessette, Christian Baron, Simon Apte, Jillian F. Banfield, Lesley A. Warren

**Author notes:** Corresponding authors: Lesley Warren,; Jillian F. Banfield. These authors contributed equally (Kelly J. Whaley-Martin, Lin-Xing Chen).

## Abstract

Acidification of freshwater in mining impacted areas is a major global environmental problem catalyzed by sulfur-oxidizing bacteria (SOB). To date, little is known about the active bacteria in mine tailings impoundments and their environmental niches. Here, biological sulfur oxidation was investigated over four years in a mine tailings impoundment, integrating sulfur geochemistry, genome-resolved metagenomics and metatranscriptomics. We demonstrated oxygen driven niche partitioning of SOB and their metabolic pathways that explain acidity generation and thiosulfate persistence. Four chemolithoautotrophic SOB, *Halothiobacillus, Thiobacillus, Sulfuricurvum* and *Sediminibacterium* comprised 37% to 73% of the analyzed communities. The impoundment waters alternated between the dominance of *Halothiobacillus* versus a *Thiobacillus, Halothiobacillus, Sulfuricurvum* and *Sediminibacterium* consortia. *Halothiobacillus* dominance was associated with lower pH values (∼4.3), higher [H^+^]/[SO_4_^2-^] and lower [S_2_O_3_^2-^], collectively indicative of extensive sulfur oxidation. *Halothiobacillus*, which couple sulfur oxidation via the Sox pathway to aerobic respiration or NO_2_^-^ reduction, were present throughout the depth profile, yet their expression of *sox* genes occurred only in upper highly oxygenated waters. Conversely, when consortia of *Thiobacillus, Halothiobacillus, Sulfuricurvum* and *Sediminibacterium* dominated, recycling/disproportionating reactions were more prevalent. *Thiobacillus,* which dominated deeper micro-oxic/anoxic waters, oxidized sulfur primarily through the rDSR pathway, coupled to NO_3_^-^/NO_2_^-^ reduction, resulting in lower [H^+^]/[SO_4_^2-^] and higher [S_2_O_3_^2-^] relative to upper waters. These field results mirror the Sox/rDSR-geochemical patterns of experimental SOB enrichments and reveal opportunities for biological treatments of recalcitrant reduced sulfur compounds, as well as gene-based monitoring and *in situ* RNA detection to predict the onset of problematic geochemistry.

## Introduction

Microbial sulfur cycling is highly complex, involving a wide range of sulfur compounds with varying stabilities, oxidation states and interlinked microbial reactions ^1–3^. Biological sulfur oxidation processes are environmentally significant as they can drive acidity generation and oxygen consumption, and are the drivers of significant environmental impacts occurring at operational mines, at legacy mining sites and in regions where sulfide-rich rocks are exposed at the surface. However, information is currently lacking on the prevalence of sulfur oxidation pathways that drive or limit these processes. Combined geochemical and “-omic” approaches have been applied to elucidate biogeochemical sulfur cycling in natural contexts, for example, in hydrothermal vents/marine settings ^4–7^, and acid mine drainage (AMD) sites ^8–10^. Lakes and man-made basins that are used as mine tailing impoundments have not been well studied with some exceptions ^11–15^. Mine tailing impoundments provide natural laboratories to examine biologically driven sulfur oxidation processes. Significantly, when sulfur oxidation intermediate compounds, industry termed “thiosalts”, discharge from impoundment waters to the natural environment, there is the potential for problematic biogeochemical reactions causing significant risk to the downstream ecosystems. Mine tailing impoundments share similarities with both natural ecosystems and AMD situations, being typically circumneutral with high metals concentrations, varied organic carbon distributions (reflecting processing reagents and natural water inputs), dynamic water geochemistry and commonly, an abundance of reduced aqueous sulfur compounds.

Previous research has identified a number of sulfur-oxidation pathways that are likely to occur in environmental settings ^16^, but their distribution and environmental factors driving selection are not well defined. Genomic analysis suggests that environmental sulfur oxidation is likely to be carried out primarily through sulfur oxidation (Sox) enzymes and the reverse dissimilatory sulfite reductase (rDSR) pathways ^16–18^, as well as the tetrathionate intermediate (S4I) pathway ^1,19^ and the Hdr pathway ^16,20–23^. The Sox pathway is well characterized and carried out by bacteria and archaea that express enzymes of the Sox complex (SoxABCDXYZ). No free reduced sulfur intermediates are produced, as they are covalently attached to the carrier proteins SoxYZ before being fully oxidized to SO_4_^2- 24^. Diverse taxonomic groups have been shown to possess Sox genes, as the Sox pathway sustains their metabolism across a wide range of geochemical conditions in the presence of reduced sulfur compounds ^16,17,25^. The rDSR pathway comprises the same proteins as the dissimilatory sulfite reduction pathway (Apr, Sat, DsrAB), while working in the reverse direction. The distribution of bacteria possessing rDSR has been documented in some environmental settings ^26^. The rDSR pathway is capable of producing free sulfur oxidation intermediates ^27^ and has been hypothesized to be more prevalent under anoxic conditions due to higher energy conservation efficiency ^28^. The intermediate tetrathionate (S4I) pathway involves the formation of free tetrathionate following the oxidation of thiosulfate catalyzed by the TsdA protein ^29^. Thus, the geochemical outcomes in nature for acidity generation and reduced sulfur compound distributions are likely to differ for each of the Sox, rDSR and S4I pathways. This is especially pertinent if these pathways are spatially or temporally segregated by differing environmental conditions such as oxygen levels. Autotrophic sulfur oxidizing bacteria that express these pathways may also have the capacity for aerobic respiration and/or denitrification, thus O_2_, NO_3_^-^ and other oxidized N compounds are plausible terminal electron acceptors for Sox, rDSR and S4I pathways. In circumneutral S-rich environmental settings where multiple terminal electron acceptors are present (i.e. O_2_, NO_3_^-^, NO_2_^-^), microbial communities may utilize an array of reactions to oxidize and/or disproportionate sulfur compounds ^30,31^, and thus ecological niche partitioning across geochemical gradients is possible.

Emerging results have identified novel sulfur-cycling organisms in circumneutral mining impacted wastewaters ^15 32^, divergent from those found in AMD and other natural sulfur rich systems ^11,12,33,34^. Mine tailing impoundments associated with sulfide ore tailings provide unique opportunities to examine microbial sulfur cycling and identify associated implications for water quality outcomes. These waters commonly possess steep oxygen gradients, multiple sulfur substrates, often present at high concentrations, as well as well adapted sulfur oxidizing microorganisms. Their investigation, therefore, also allows us to test hypotheses concerning possible sulfur pathway niche partitioning in a highly dynamic, environmental setting. In this field study, depth dependent physico-chemistry, S, N and organic carbon geochemistry, microbial community structure and functional metabolic capabilities associated with a tailings impoundment water column at an active Ni/Cu mine site (Figure 1) were analyzed over four years (2015-2018). We discovered that microbial sulfur metabolic strategies are spatially and temporally segregated by oxygen availability and that this niche partitioning, particularly with regards to *Halothiobacillus*, determined the initiation and extent of acidity generation.

**Figure 1.**
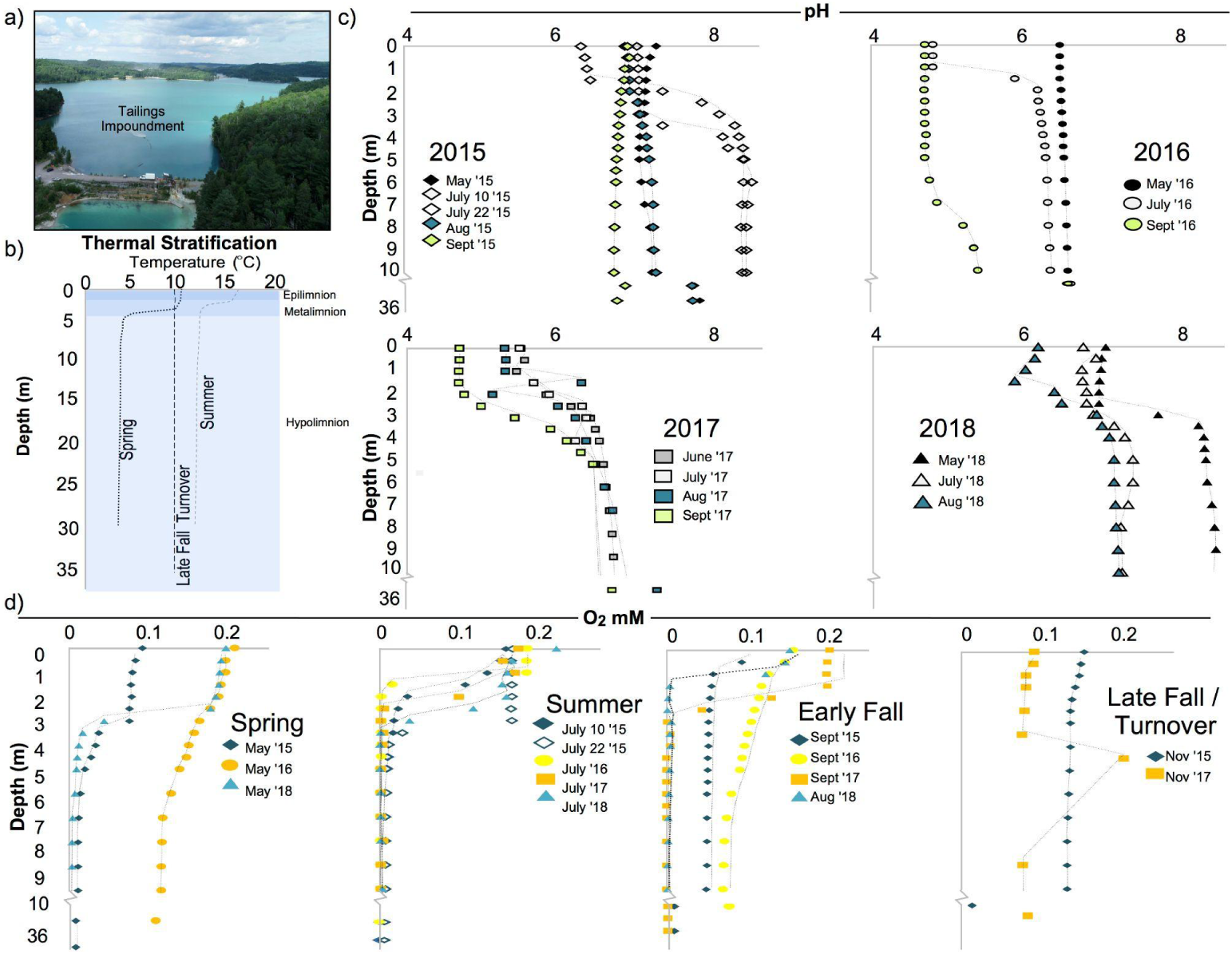
Physiochemical depth profiles in tailing impoundment waters from 2015 to 2018 (0-10 m; comprising the epilimnion, metalimnion and upper hypolimnion when stratified, of the ∼38 m water column). (a) Drone photograph of tailings impoundment b) Representative seasonal temperature profiles (Spring, Summer, Late Fall Turnover) for the dimictic tailings impoundment (c) Available monthly pH depth profiles by year: 2015, 2016, 2017 and 2018. (d) Oxygen (mM) depth profiles by season across years: Spring (May), Summer (July), Early Fall (August, September) and Late Fall turnover (November).

## Materials and methods

### Site description

The tailings impoundment is the central repository for mining impacted water and sulfidic mineral tailings from active local mining operations with a total volume of 19,975,000 m^3^ and a maximum depth of ∼38 m (Figure 1). This systems was designed from a pre-existing lake to inhibit the oxidation of the deposited high sulfide tailings, yet encourage oxidation of aqueous surficial sulfur oxyanions (e.g., S_2_O_3_^2-^, S_3_O_6_^2-^, S_4_O_6_ ^2-^) to sulfate (the mill commonly discharges inputs with higher concentrations of SO_3_ ^-^ and S_3_ O_3_ ^2-^) prior to discharge into a river system and local watershed ^35^. This basin forms the beginning of a large-scale water treatment system at a Ni/Cu mine in Sudbury, Ontario, Canada ^13,15^ with dynamic inputs from various sources including ore processing waters, tailings deposition and natural tributaries. The impoundment is dimictic, with summer (open water) and winter (under ice) thermal stratification periods with residence times estimated in the epilimnetic portions of the water column to be on the order of months.

### Water sampling and processing

During 21 sampling campaigns, water samples were collected from the impoundment at various depths over four years from March 2015 to August 2018 during mostly open water periods (i.e., no ice cover; with the exception of March 2015; Table S1). The sampling procedures have been described in detail elsewhere ^15,36^. Briefly, water samples were collected from identified depths during each campaign from a floating platform accessible by boat in ∼38 meters of water cap depth in the impoundment and if unavailable, from shore locations. Prior to sampling, a physico-chemical profile of the impoundment water was collected using a multiprobe (temperature, O_2_, pH, ORP, conductivity; YSI 600 XLM). Physicochemical survey data for the impoundment water column for all 21 sampling campaigns are provided in Table S1. Water samples were collected by a VanDorn water sampling bottle (Wildco, 3.2 L Horizontal beta™) during stratified periods from the epilimnion, metalimnion and hypolimnion, and from upper and lower depths during mixing periods. In addition, during some campaigns surface water samples (0 - 0.5 m) were also collected from shore utilizing a telescopic sampler. Water samples were collected for geochemical analysis (n=53; sulfur and nitrogen speciation, dissolved organic carbon (DOC)) and for molecular biology analysis (n=30; 16S rRNA gene sequencing and metagenomics as well as metatranscriptomics (RNA) analysis in August 2018 (n=3). A subset of these *in situ* waters were taken through laboratory sulfur oxidative enrichment experiments along with wastewaters collected from three other mine sites in Canada (Flin Flon, Manitoba; Snow Lake, Manitoba; Baie Verte, Newfoundland). Detailed methods, geochemical and 16S rRNA gene sequencing results were described previously ^15^. Here, we present the metagenomic results of those sulfur oxidizing enrichments to align with the field-based metagenomes.

Water samples for SO_4_^2-^ and nitrogen species (NH_4_^+^, NO_2_^-^, NO_3_^-^) quantification were collected in 250 ml to 1 litre polypropylene bottles (each rinsed three times with site water prior to collection) with no headspace and stored at 4°C prior to analysis (within 24-72 hours). ΣH_2_S was determined on-site rapidly after sample collection as previously described ^36^. Thiosulfate (S_2_O_3_^2-^) and sulfite (SO_3_^2-^) analytes were stabilized immediately upon sampling through derivatization with monobromobimane to prevent oxidation ^37^. In detail, water samples (100 μL) were directly pipetted into 2 mL glass amber vials with derivatization agents (100 μL of acetonitrile, 100 μL of 50 mmol L^-1^ HEPES (4-(2-hydroxyethyl) piperazine-1-ethanesulfonic acid, ≥99.5%, Sigma), 5 mmol L^-1^ of EDTA buffer (ethylenediaminetetraacetic acid, 99.4-100.6%, Sigma Aldrich; pH=8.0, adjusted with NaOH) and 20 μL of 48 mmol/L monobromobimane (>97%, Sigma Aldrich). After 30 minutes of reaction time, 200 μL of methanesulfonic acid (∼100 mmol) was added (≥99.5%, Sigma Aldrich) and samples were subsequently kept frozen until analysis. For total sulfur (TotS) analyses, water samples were collected in triplicate (40 mL) *in situ* and filtered through Pall Acrodisc^®^ 25mm 0.45 μm Supor^®^ membrane filters with polypropylene syringes) into 50 Falcon™ tubes pre-spiked, to achieve final concentrations of 0.2% HNO_3_ (Optima grade, Fisher Chemical) and then stored at 4°C until further processing. Water samples for TOC and DOC concentrations were collected in acid washed amber glass bottles pre-combusted at 450°C for 8 hours and frozen until analysis. Samples for DNA and RNA analyses were filtered (∼2 to 5 L) through triplicate 0.2 and 0.1 μm filter towers (Thermo Scientific™ Nalgene™ Rapid-Flow™ Sterile Disposable Filter Units with CN Membrane) until filters clogged. Filters were excised and kept frozen (−20°C for DNA, -80ºC for RNA) until nucleic acid extraction.

### Genomic DNA extraction and quantification

The DNA extraction was conducted on filters using DNeasy PowerWater DNA Isolation Kit (QIAGEN), and isolation was carried out with recommended protocols (QIAGEN). DNA extracts were sent to the Farncombe Metagenomics Facility at McMaster University, (Hamilton, Ontario) for subsequent analysis. Library concentrations were quantified through quantitative PCR (qPCR) and concentrations were adjusted for each step of the subsequent molecular protocol.

### RNA extraction, library generation and sequencing

The extraction of total RNA was conducted on filters using the RNeasy PowerWater Kit (from Qiagen). The concentration of total RNA was determined using a Nanodrop instrument and the quality of the preparation was assessed by agarose gel electrophoresis to monitor 16S and 23S ribosomal RNA. Samples were stored at -80°C; samples from two extractions were pooled before library construction. Quality controls, rRNA depletion, cDNA library construction from isolated RNA and sequencing were performed at the Génome Québec Innovation Centre (Montréal, Canada). Sequencing of the libraries was done using Illumina NovaSeq technology (NovaSeq 6000 S4, 100 bases paired-end). Sequences obtained in this study were deposited at NCBI under the bioproject accession number PRJNA763576.

### Barcoded 16S rRNA gene amplicon sequencing and bioinformatics analyses

Aliquots of purified DNA were used to amplify region V4 of the 16S rRNA gene by PCR using Illumina adapted primers following Bartram et al.^38^ and standard protocols of the Earth Microbiome Project ^39,40^. In short, the primers were modified to amplify 515f (GTGYCAGCMGCCGCGGTAA) and 806r (GGACTACNVGGGTWTCTAAT) variable regions of archaeal and bacterial 16S rRNA gene. PCR was performed using 50 ng of the template and the PCR mix contained 1U of recombinant Taq DNA Polymerase (Invitrogen™), 1x buffer, 1.5 mmol L^-1^ MgCl2, 0.4 mg mL^-1^ BSA, 0.2 mmol L^-1^ dNTPs, and 5 pM of each primer. The reaction was carried out at 98°C for 5 minutes, 35 cycles (98°C) for 30 seconds, then 30 seconds (50°C) and 30 seconds (72°C), with a final extension of 72°C for 10 minutes. PCR products were checked by electrophoresis and sent for sequencing. All amplicons were normalized to 1.25 ng μL^-1^ using the SequalPrep normalization kit (ThermoFisher#A1051001) and sequenced using the Illumina MiSeq platform. The percentage of reads that passed the quality control parameter “Q30” (error rate = 0.1%) for each of the triplicates runs were 85%, 89% and 90% respectively. Bimera checking was performed on the data carried out through DADA2 (version 1.6.0) and 3-5% of reads were detected as bimeras were excluded ^15^. Raw sequences were filtered and trimmed with a minimum quality score of 30 and a minimum read length of 100bp using Cutadapt ^41,42^. DADA2 version 1.6.0 ^42^ was utilized to resolve sequence variants. Bimeras, and chloroplast and mitochondrial sequences were identified and removed. SILVA database version 132 ^43^ was used to assign a taxonomy based on 16S rRNA sequences.

### Metagenomic sequencing, reads processing and assembly

The genomic DNA samples extracted from filters (duplicates of those used for 16S rRNA gene sequencing analyses) were also used for metagenomic sequencing. Preserved DNA extracts (n=9) from sulfur oxidation enrichments described in Whaley-Martin et al.^15^ were also subjected to metagenomic sequencing. The DNA extracts were dried and resuspended in 25μL of water. Construction of libraries (insert length ∼ 500 bp) and sequencing by Illumina HiSeq 1500 with paired-end 150 bp sequencing kit, were performed at the Farncombe Metagenomics Facility at McMaster University. The raw paired-end reads of the metagenomic sequencing samples were filtered to remove Illumina adapters, PhiX and other Illumina trace contaminants with BBTools ^44^, and low quality bases and reads using Sickle (version 1.33). The *de novo* assembly was performed with IDBA_UD ^45^ (parameters: --mink = 20, --maxk 140, --step 20, --pre_correction). In order to determine the sequencing coverage of each scaffold from a given sample, quality reads sets from all samples were individually mapped to the full assembly using Bowtie2 ^46^ with default parameters. The sam file was converted to bam format and sorted using samtools ^47^, and the coverage of each scaffold from a given sample across all samples was calculated using the *jgi_summarize_bam_contig_depths* script from MetaBAT ^48^. All the scaffolds with a minimum length of 2500 bp were assigned to genomes bins using MetaBAT ^48^, with both the tetranucleotide frequency and sequencing coverage across all samples considered. All the binned and unbinned scaffolds with a minimum length of 1000 bp were uploaded to ggKbase (ggkbase.berkeley.edu) for manual genome curation, based on GC content, sequencing coverage and taxonomic information of each scaffold as previously evaluated ^49^.

### Gene prediction and annotation

The assembled scaffolds with a minimum length of 1 kbp (hereafter “1k_scaffolds”) were included for gene prediction and subsequent analyses. The protein-coding genes were predicted by Prodigal V2.6.3 ^50^ from 1k_scaffolds (parameters: -m -p meta). The 16S rRNA genes were predicted from 1k_scaffolds based on an HMM database that reported previously ^51^. The tRNAs encoded on all 1k_scaffolds were predicted using tRNAscanSE (version 2.0.3) ^52^. The predicted protein-coding genes were compared against the databases of Kyoto Encyclopedia of Genes and Genomes (KEGG) ^53^, UniRef100 ^54^ and UniProt ^55^ using Usearch (version v10.0.240_i86linux64) ^56^ for annotation. For specific metabolic potentials that are of interest, the predicted protein-coding genes were also searched against the HMM databases reported previously ^57^.

### Genome comparison

To identify the similarity of the genomes for each genus that is of interest, the “compare” function of dRep ^58^ was used on the genomes modified by ggKbase (ggkbase.berkeley.edu).

### RNA-Seq analysis

The raw RNA sequencing reads were conducted for quality control as performed for metagenomic reads (see above). The quality RNA reads were mapped to the nucleotide sequences of protein-coding genes from the corresponding samples, using Bowtie2 ^59^ with default parameters. The transcriptional level of each protein-coding gene was determined by calculating its RNA sequencing coverage using the *jgi_summarize_bam_contig_depths* script from MetaBAT ^48^.

### Relative abundance calculation of functional genes

To calculate the relative abundance of each protein-coding gene of interest, the total coverage of all genes in each sample was firstly determined as: Total_cov_ = *G*_*ij*_ x C_*j*_, where *G*_*ij*_ is the number (i.e., *i*) of protein-coding genes on scaffold *j*, and C_*j*_ is the sequencing coverage of scaffold *j* (see above). The coverage of a specific gene (for example, gene A) is determined as: Gene_A_cov_ = C_*j*_, where C_*j*_ is the sequencing coverage of scaffold *j* that encodes gene A. Thus, the relative abundance of gene A in the sample is determined as below: Rel_abun_A_ = C_*j*_ x 100 / Total_cov_ %.

### Phylogenetic analyses

To distinguish the function (oxidation or reduction) of the detected *dsrAB* genes, a phylogenetic tree was built for concatenated DsrA and DsrB protein sequences, with references from Anantharaman et al. ^26^. The DsrA and DsrB sequence sets were respectively aligned with MUSCLE ^60^ and filtered using trimAl ^61^ to remove those columns with at least 90% gaps, then concatenated using Geneious Prime ^62^ in the order of DsrA-DsrB. The phylogenetic tree was built using IQtree ^63^ with 1000 bootstraps and the “LG+G4” model. The oxidative type of DsrAB was determined according to Anantharaman et al. ^26^.

### Sulfur and nitrogen speciation analyses

Dissolved ΣH_2_S, SO_4_^2-^, NH_4_^+^, NO_2_^-^, NO_3_^-^ concentrations were determined on unfiltered samples by spectrophotometry using a HACH DR2800 (HACH Company, Loveland, CO, USA) ^15,36,64^. Thiosulfate and sulfite concentration analyses (field stabilised monobromobimane derivatives) were carried out with methods adapted from ^37^ on a Shimadzu LC-20AD prominence liquid chromatography (LC) system coupled to a fluorescence UV/VIS detector on an Alltima™ HP C18 reversed phase column (150 mm × 4.6 mm × 5 μm, Grace™) held at 35°C using an isocratic mobile phase of 65% of 0.25% acetic acid v/v (pH 3.5 adjusted with NaOH) and 35% methanol at 0.5 mL min^-1^ for a total run time of 12 minutes as described previously^15^. The excitation wavelength was set to 380 nm and the emission wavelength 478 nm. Sulfite eluted at ∼5.1 minutes and thiosulfate eluted at ∼5.5 minutes. Calibration curves were prepared using sodium sulfite (Sigma Aldrich, ≥98% purity) and sodium thiosulfate (Sigma Aldrich, 99% purity). All sulfur compound concentrations reported represent the sulfur molarity (i.e. S in S_2_O_3_^2-^).

### Total sulfur (TotS) analysis

Total sulfur concentration analyses were conducted at the Commonwealth Scientific and Industrial Research Organization (CSIRO), Sydney, Australia by inductively coupled plasma optical emission spectroscopy (Varian730 ES, Mulgrave, Australia). Varian Fast Automated Curve-fitting Technique (FACT) was used to correct for background/inter-element interferences. Calibration standards in 2% v/v HNO_3_ were prepared from certified reference stocks (AccuStandard New Haven, CT, USA). The limit of detection (LOD) for TotS was 1 mg/L.

### Total and dissolved organic carbon analyses

Frozen samples were thawed and for DOC analysis a sub-sample was filtered through Pall Acrodisc^®^ 25mm 0.45 μm Supor^®^ membrane filters using polypropylene syringes and placed into carbon clean 40mL glass vials. Organic carbon concentrations were determined on a Shimadzu TOC-L using a 680°C combustion and non-dispersive infrared (NDIR) detection. Inorganic carbon samples were acidified (25% H_3_PO_4_) and sparged prior to combustion analysis using the inorganic carbon (IC) reactor kit attachment. Calibration standards were prepared using potassium hydrogen phthalate (Sigma Aldrich, 99.95-100.05% purity) for DOC, sodium hydrogen carbonate (Sigma Aldrich, ≥99.7% purity) and sodium carbonate (Sigma Aldrich, ≥99.5% purity) for IC standards (limits of detection for DOC were 1 mg/L C). DOC values were determined using subtraction of IC values from total carbon concentrations for each filter fraction.

### Statistical and data analysis

Non Metric Dimensional Scaling (NMDS), Bray-Curtis Dissimilarity Clustering and Pearson R correlation matrices were carried out in R version 1.1.463 utilizing the Vegan package version 2.4-5. Chemical and biological data where concentrations/abundances were less than the limits of detection were treated as zero for statistical analysis.

## Results

### Water column depth profiles of pH and oxygen show steep gradients and seasonal variations

The pH depth profiles showed circumneutral conditions throughout the water column in 2015 and 2018 (ranging from ∼8.6 - 6.2), while 2016 and 2017 impoundment waters were more acidic (ranging from ∼ 6.6 - 4.7) (Figure 1c). The lowest pH values were observed in upper waters during the summer months of 2016 and 2017 (Figure 1c). Dissolved oxygen concentrations in May 2016 (ranging from 0.11 to 0.21 mM; data for May 2017 not available) were considerably higher than those observed in May 2015 and 2018 (ranging from 0.006 to 0.09 mM) (Figure 1d). July oxygen profiles across all years showed the establishment of steep oxygen gradients in the summer thermally stratified water cap, with epilimnetic upper waters (∼0-2 m) oxygen concentrations ranging from 0.17 to 0.24 mM and lower metalimnetic and hypolimnetic waters (> 3 m depth to 38 m) oxygen concentrations ranging from 0.03 mM to < limit of detection (LOD). In September of 2015 and 2016 (prior to fall turnover), a rebound in dissolved oxygen concentration occurred after a planned summer mine shut-down event, where sub-aqueous tailing deposition ceased for a two-week period suggesting the perturbation caused a decrease in oxygen consuming processes. Lake turnover events in November 2015 and 2017 coincided with a more uniform/mixed oxygen profile throughout the water column (Figure 1d).

### Temporal trends in acidity generation and sulfur distributions correlate with pH values

The H^+^/SO_4_^2-^ ratio provides a comparative indicator of overall acidity generation associated with microbial sulfur oxidation metabolism ^65^. Higher values indicate oxidation of reduced sulfur compounds to sulfate accompanied by higher acidity generation, whereas lower values are indicative of more complex sulfur recycling/disproportionation (and often acid consuming) reactions. Comparison of H^+^/SO_4_^2-^ values revealed significantly higher values in samples from 2016 compared to any of the other three years (ANOVA and a post-hoc Tukey comparison test; p < 0.05) (Figure 2a). H^+^/SO_4_^2-^ values were also elevated in 2017, which is consistent with the lower pH values observed in 2016/2017. Thiosulfate concentrations in 2018 were statistically higher (p < 0.01) than those measured in 2016 and 2017, but were not statistically significantly different from 2015 thiosulfate concentrations (ANOVA and a post-hoc Tukey comparison test; p < 0.01) (Figure 2b). Total sulfur (TotS) (Table S2) increased over the four years of investigation in the summer waters. [SO_4_^2-^] and [reactive S] (S_react_ = TotS – SO_4_^2-^, i.e., all possible S atoms that can take part in oxidative or disproportionation reactions; ^36^ were variable over time showing no evident temporal trends. Sulfide (∑H_2_S) concentrations ranged from < 1 μM to 6 μM with progressive increase with depth.

**Figure 2.**
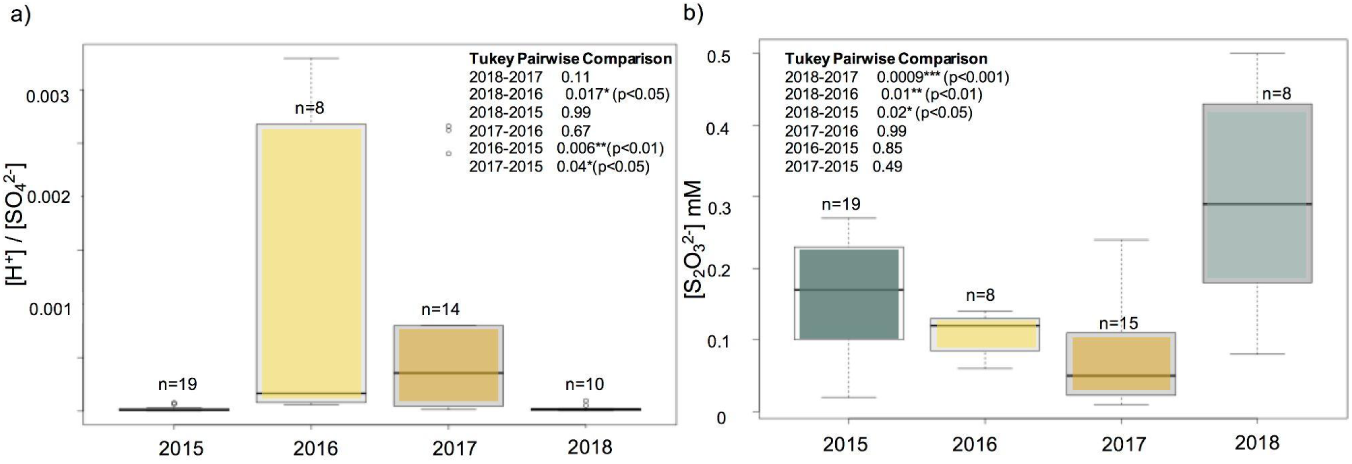
Sulfur indicators of various sulfur oxidative microbial pathways across time (2015, 2016, 2017 and 2018). (a) Box-plot and statistical (ANOVA and subsequent Tukey pairwise comparison) of annual sulfur speciation ratios of [H^+^]/ [SO_4_^2-^] (omitting May 2017 0.5 m and May 2018 21 m where both pH and SO_4_^2-^ were unavailable), and (b) Annual observed [S_2_O_3_^2-^] (mM) in tailings impoundment waters inclusive of March 2015 to August 2018 with the exception of May 2018 (0.5 m), July 11, 2018 (2.8 m) July 23, 2018 (0.5 m) when data were not available. Two outliers included in the statistical comparison were excluded from the graph for visualization purposes (March 2015 at 21 m (0.59 mM) and June 2018 at 0.5 m (1.7 mM); see Table S4).

### Microbial community composition correlates with acidity generation

A total of 3,533,030 16S rRNA gene sequences were obtained from 30 water samples collected throughout 2015-2018. Overall, average Shannon Diversity Indices (H’) decreased from 2.5 in 2015 to 1.9 for 2016/2017 and increased to 3 in 2018 (Table S3). Both 16S rRNA gene based (Table S4) and metagenomic analyses (Table S5) indicated that these waters were inhabited by communities that varied across time in composition and diversity, but were dominated by chemolithoautotrophic sulfur oxidizing bacteria over all 4 years (∼76% in 2015, ∼55% in 2016/2017 and ∼60% in 2018) (Figure 3a, Table S4).

**Figure 3.**
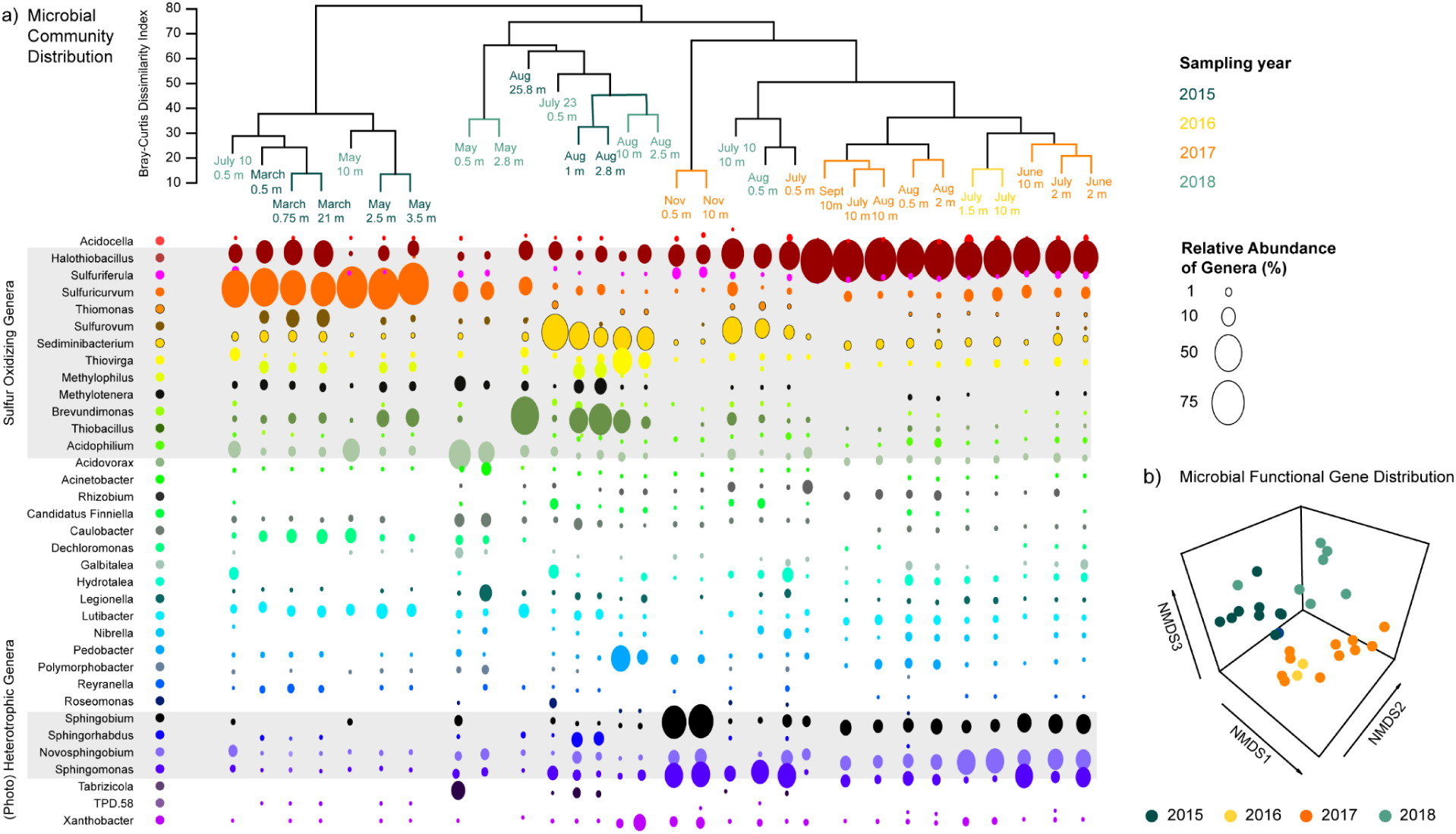
Microbial community composition over time. (a) Bray Curtis dissimilarity clustering of microbial communities and bubble plot representation of relative abundances of bacterial genera (showing only genera present at > 1% within at least two samples). (b) Non-metric dimensional scaling (NMDS) (k=3) representation of the microbial community functionality (derived the relative abundance of sulfur cycling, nitrogen cycling, hydrogen cycling, carbon fixation, photosynthetic and aerobic respiration genes (Table Sx) (n=30) Stress = 0.065, non-metric fit = 0.996, linear fit, R^2^ = 0.97).

Overall, *Halothiobacillus, Sulfuricurvum, Thiobacillus* and *Sediminibacterium* were the most abundant genera. *Halothiobacillus* was the dominant sulfur bacterium in 2016 and 2017 and this correlated with lower overall diversity and acidification. In contrast, significantly higher proportions of *Sulfuricurvum, Thiobacillus* and *Sediminibacterium* were detected in 2015 and 2018 water samples, correlating with lower abundance of *Halothiobacillus sp.* (Figure 3a). Specifically, the tailing impoundment waters from 2015 (n=8) were dominated by chemolithoautotrophic sulfur oxidizers/disproportionating *Sulfuricurvum* (37 ± 28 %), *Thiobacillus* (17 ± 20 %), *Halothiobacillus* (12 ± 8 %), and *Sediminibacterium* (5.9 ± 9 %) (Table S4). The primary chemolithoautotrophic sulfur bacterium in 2016 and 2017 was *Halothiobacillus sp.* (50 ± 21 %), while *Sulfuricurvum* and *Sediminibacterium* was much less present (total ∼3%), and *Thiobacillus* was undetectable (Figure 3a, Table S4). In 2018, there was a similar pattern to 2015 with all four dominant genera detected, showing a rebound in *Sulfuricurvum* (17 ± 26 %) and a re-emergence of *Thiobacillus* (3.9 ± 6 %), while *Halothiobacillus* abundances decreased (11 ± 11 %) (Figure 3a, Table S4). Greater diversity was observed in the microbial communities from 2018, indicated by the presence of all four of these major sulfur metabolizing groups, as well as *Thiovirga* (5 ± 8 %) and *Acidovorax* (9 ± 11 %) and other non-sulfur microorganisms (Table S4). Two major shifts in the microbial community coincided with major mine operational changes during July of 2015 and 2018 (“shutdown events” where sub-aqueous tailing deposition ceased for a two-week period in July 2015 and 2018); in both years the abundance of *Sediminbacterium* and *Thiobacillus* increased (Figure 3a).

Bray-Curtis Cluster and non-metric-dimensional scaling (NMDS) 3-dimensional analysis revealed three temporal clusters for both microbial community structure and metabolic capability (inferred by gene relative abundance; Table S5 (Figure 3c), and those from 2016 and 2017 were more similar compared to 2015 or 2018. These community structure clusters were driven by abundance changes for *Halothiobacillus sp.* (dominance in 2016/2017) relative to higher abundances of *Sulfuricurvum, Thiobacillus* and *Sediminibacterium* in 2015 and 2018 (Figure 3a).

Regardless of which chemolithoautotrophic sulfur cycling genera were present across the four years, they always co-existed with known (photo)-heterotrophic organisms that have been reported to be metal tolerant ^66,67^, including the families Flavobacteriaceae (*Lutibacter*) and Sphingomonadaceae (*Sphingomonas, Novosphingobium, Sphingobium*) (Figure 2a, Table S3). In 2016 and 2017, *Halothiobacillus sp.* and members of the *Sphingomonadaceae* family (heterotrophs; *Sphingomonas, Novosphingobium* and *Sphingobium*) dominated the tailing impoundment waters, comprising an average of 31 % of the communities (Table S4). In 2016/2017 waters there was a progressive decrease in the (photo)-heterotrophic family *Sphingomonadaceae* and increase in *Halothiobacillus* with depth (Pearson test: R = - 0.93, p = < 0.0001) (Figure S2). A slight decrease in DOC concentrations occurred in 2016 and 2017 when heterotrophic bacteria were higher in abundance (Table S6).

### The distribution of sulfur oxidation genes at temporal and spatial scales

At the community level over the four years of the study, *soxABCDXZ* complex genes were consistently present while *dsrABCEFH, sat and aprAB* were limited to 2015 and 2018 (Figure 4a; Tables S5). Notably, DsrC was not distinguishable in the HMM database from other bacterial proteins that belong to the TusE/DsrC/DsvC family. Thus, while DsrC is considered a key protein the rDSR sulfur oxidation pathways and abundances are reported (Tables S5), the total abundance of DsrC was not represented in Figure 4a ^68^. TsdA was also consistently present across time and space, while *ttrABC* showed a similar trend to the *rDSR* gene that was only present in 2015 and 2018 (Figure 4a;b). A direct comparison of gene distribution with [H^+^]\[SO_4_^2-^] ratios and [S_2_O_3_^2-^] concentrations (n=30) (Figure 4a) revealed a relationship between overall sulfur geochemical outcomes and sulfur gene distributions. The 2015 and 2018 waters hosted microbial communities with functional S genes encoding the Sox (SoxABCDXYZ), S4I (TsdA) and rDSR (DsrABCEFH, AprAB and Sat) pathways, lower acidity to sulfate ratios (driven by similar dissolved sulfate concentrations but significantly greater acidity) and higher thiosulfate concentrations. Geochemical monitoring of the input waters discharging into the tailings impoundment over the course of this study revealed (data not shown) no discernible change in overall acidity loading over the course of this study. These results are suggestive of greater overall sulfur recycling/disproportionating and/or slower oxidative reactions occurring in 2015 and 2018 relative to 2016 and 2017 (Figure 4a). In 2016 and 2017 when the highest acidity to sulfate ratios and lowest thiosulfate concentrations occurred, the most abundant S genes encoded the Sox (SoxABCDXYZ) and S4I (TsdA) pathways. The geochemical trends of this comparison were consistent with patterns observed from the larger geochemical dataset (H^+^/SO_4_^2-^, n=51; and S_2_O_3_^2-^concentrations, n=50) (Figure 2). In 2015 and 2018, when *Sulfuricurvum, Thiobacillus* and *Sediminibacterium* were dominant, the abundances of genes encoding the rDSR pathway were inversely correlated with oxygen (Figure S3). *DsrABCEFH, aprAB* and *sat* genes (possessed by *Thiobacillus* spp) (Figure 5) showed greater abundance within the micro-oxic to non-detectable oxygen (< LOD to 0.003 mM) portions of the water column (Figure S3) in March 2015, August 2015 and August 2018.

**Figure 4.**
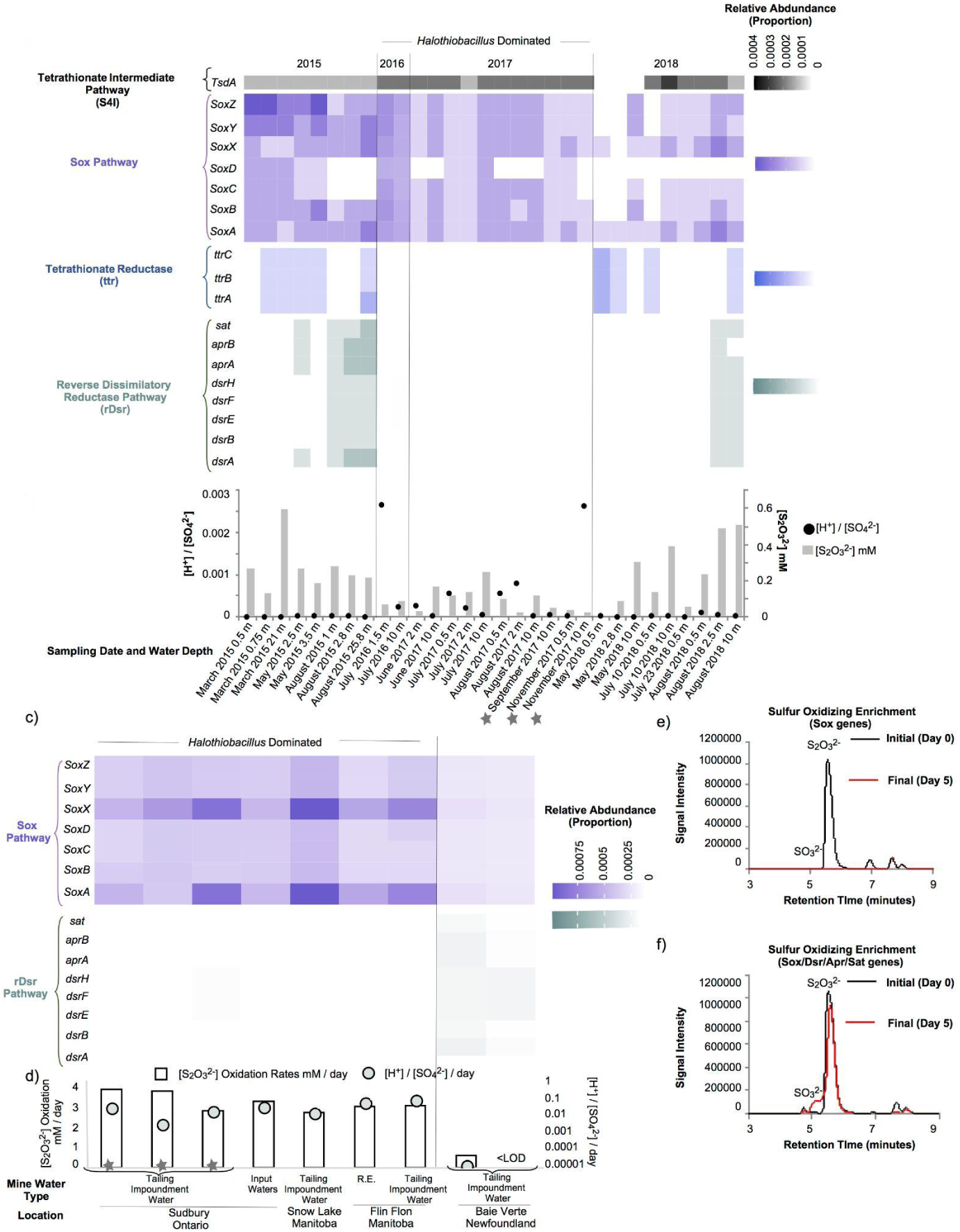
Dissimilatory sulfur gene distributions and sulfur geochemistry of natural tailings impoundment waters and cross-mine laboratory enrichments. a) The tailings impoundment community level gene relative abundance of (a) *TsdA, soxABCDXYZ, ttrABC dsrABEFH*, a*prAB* and *sat.* (b) The distribution of [H^+^]:[SO_4_^2-^] and S_2_O_3_^2-^ concentrations from 2015 to 2018 in the tailings impoundment. (c) community level gene relative abundances of sulfur oxidizing bacterial laboratory enrichment from four mine sites (*SoxABCDXYZ, dsrABEFH,* a*prAB* and *sat).* (d) the daily rate of S_2_O_3_^2-^ oxidation and rate change of [H^+^]:[SO_4_^2-^] / day, the initial and final chromatograms (HPLC-UV/Vis) showing dissolved sulfite and thiosulfate in the sulfur oxidizing bacteria enrichment waters with genes, (e) of an enrichment with functional genes *soxABCDXYZ* and (f) another enrichment with *soxABCDXYZ* and *dsrABEFH, aprAB, sat.* R.E. stands for receiving environment (water discharged from the mine treatment system into the environment and star symbols indicate the three tailing impoundment waters from Sudbury, Ontario that were also enriched for sulfur oxidizing bacteria (as described previously ^15^).

**Figure 5.**
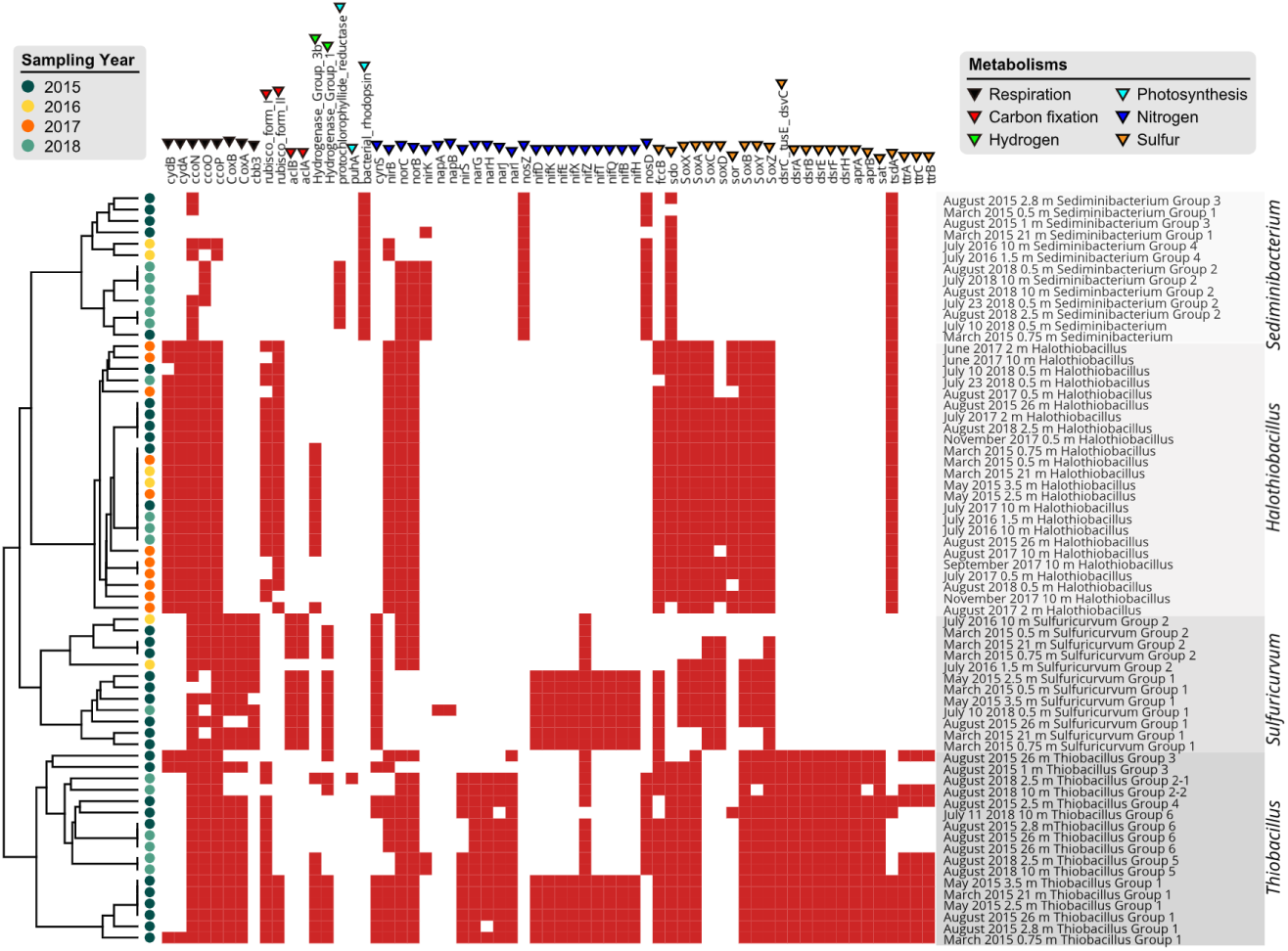
The distribution of genes in key metabolisms. The presence of a given gene in a genome is indicated by a red square, its absence by a white square. The genes are shown for the dominant sulfur oxidizing genera in the tailings impoundment waters, *Thiobacillus, Sulfuricurvum, Halothiobacillus* and *Sediminibacterium.* The functional categories of the genes are indicated by colored triangles. The genomes are clustered based on the presence and absence of listed genes.

Sulfur oxidizing bacteria enriched directly from *in situ* mine waters provide an opportunity to examine gene driven differences in sulfur oxidation rates and sulfur intermediate formation under laboratory settings. Here, metagenomic and geochemical analysis of sulfur oxidizing bacteria enrichment experiments grown from waters collected at this and three other mine tailings impoundments revealed the same connections between genera, Sox vs rDSR gene presence and sulfur outcomes as identified in our field results (Figure 4b). Bacterial enrichments that solely contained Sox genes had faster oxidation rates and did not produce detectable sulfite compared to enrichments in which rDSR genes were also present (Figure 4b). SoxAX genes were consistently the most abundant in all of the enrichments which may be due to the bacteria possessing multiple copies of these genes that has also been observed in a previous study ^69^.

### Genomic and metabolic divergences of the dominant genera

Genome-resolved metagenomics analysis was used to reconstruct genomes of bacteria from the four most dominant genera and other species with genes for sulfur-oxidizing pathways. The genomic and metabolic diversity of the dominant sulfur cycling bacteria were compared. *Halothiobacilli* comprising a single group (Figures 5 and 6) had very similar genomes (24 genomes sharing > 99% nucleotide identity) that encode the Sox complex, Sor (sulfur oxygenase reductase), Sdo (sulfur dioxygenase), TsdA (thiosulfate dehydrogenase) and FccB (flavoprotein subunit of flavocytochrome) proteins of S metabolism. In contrast, *Sulfuricurvum* comprised two distinct groups (12 genomes), all with genomes that encode the Sox complex. *Sediminibacterium* formed four groups (13 genomes) and each genome contains the *sdo* and *tsdA* genes. The greatest diversity existed within the *Thiobacillus* genus (17 genomes). Seven groups were present in the tailings impoundment in 2015 and groups 1, 3, 6, and 7 dominated while in 2018 groups 2, 4 and 5 dominated. All *Thiobacillus* groups encoded Dsr (dissimilatory sulfur reductase), Apr (adenosine 5′-phosphosulfate reductase), Sat (sulfate adenylyltransferase), Sdo and the Sox complex (Figures 5 and 6) with cross group variation evident in (by?) the presence of the *tsdA* gene. Other less abundant genera (*Brevundimonas, Methylotenera, Sulfurovum* and *Thiomonas*) possessed the capacity for sulfur oxidation through the Sox pathway (Figure 6).

**Figure 6.**
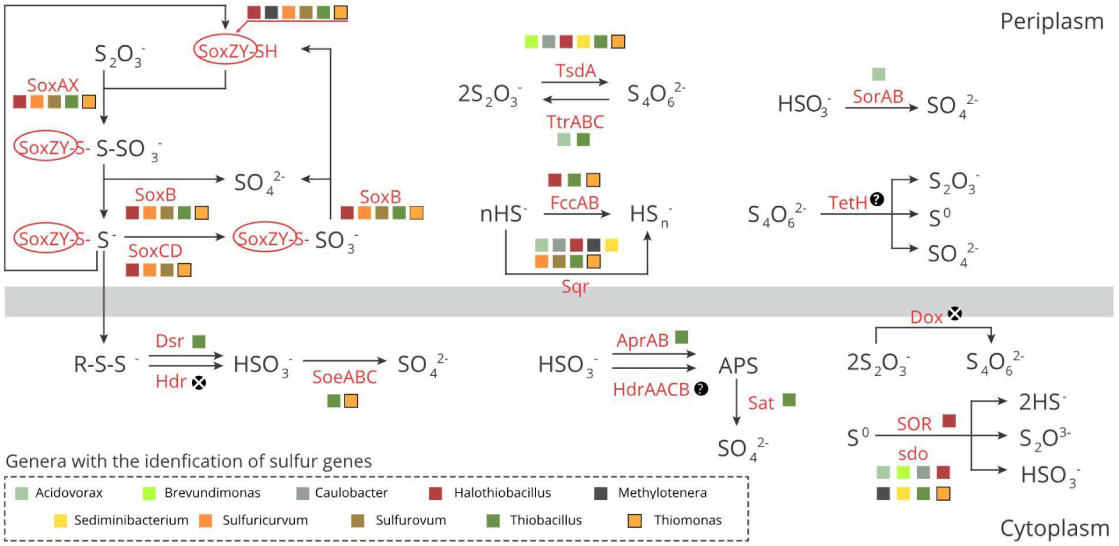
Schematic of potential sulfur oxidation pathways based on identified genes encoding sulfur metabolic enzymes. A black circle with “X” indicates the gene was not detected in the genomes of the listed genera, a black circle with “?” indicates no appropriate HMM database is available for reliable annotation. This pathway representation was adapted from Watanabe et al.^16^.

### Transcriptional activities of metabolic genes

Since the presence of genes encoding S metabolic proteins alone does not show that a pathway is being expressed, we isolated mRNA and conducted RNA-seq analysis of samples collected in the peak summer stratified period of August 2018 (depth of 0.5 m, 2.5 m and 10 m). Results showed that the patterns of gene expression of *Halothiobacillus* sp. and *Thiobacillus* spp. (“Groups 1, 4 and 5”) are consistent with the hypothesis that segregation of microbial sulfur oxidation metabolic pathways occur across this steep oxygen gradient (Figure 7). The results of RNA-Seq analysis support the expression of three sulfur oxidation pathways: Sox, rDSR and S4I. *Halothiobacillus* activity (Sox and S4I pathways) was mostly limited to the upper oxygenated epilimnetic waters, expressing aerobic respiration genes (*ccoNOP, cydAB*), and possibly, nitrite and nitric oxide reduction (*nirB* and *norBC* activity) coupled to sulfur oxidation (Figure 7c). Notably, *Halothiobacillus sox* gene expression was mostly limited to the most oxic portions of the stratified water cap (epilimnetic and upper metalimnetic regions) (Figure 7), despite its occurrence within the anoxic hypolimnetic regions (Figure 3a). In contrast, gene expression of *Thiobacillus spp*. (including genes encoding the rDSR pathway), an incomplete Sox (lack of SoxCD) and S4I (TsdA) was confined to the low oxygen (2.5 m at 0.0016 mM O_2_) metalimnion (Groups 1, 4 and 5) and upper hypolimnion (10 m at 0.0006 mM O_2_) (Groups 1 and 5) regions. In line with this position in the water column, we observed the expression of genes encoding enzymes for nitrite and nitric oxide reduction (*nirB* and *norBC*) as well as sulfur oxidation (*dsrABCEFH, sat, aprAB*) for Group 2 *Thiobacillus spp*. in the microoxic metalimnion and anoxic hypolimnion regions (Figure 7c.) Surprisingly, while *Sediminibacterium* occurred in relatively high abundance in the samples (Figure 1), their transcriptomic activity was apparently low (not shown).

**Figure 7.**
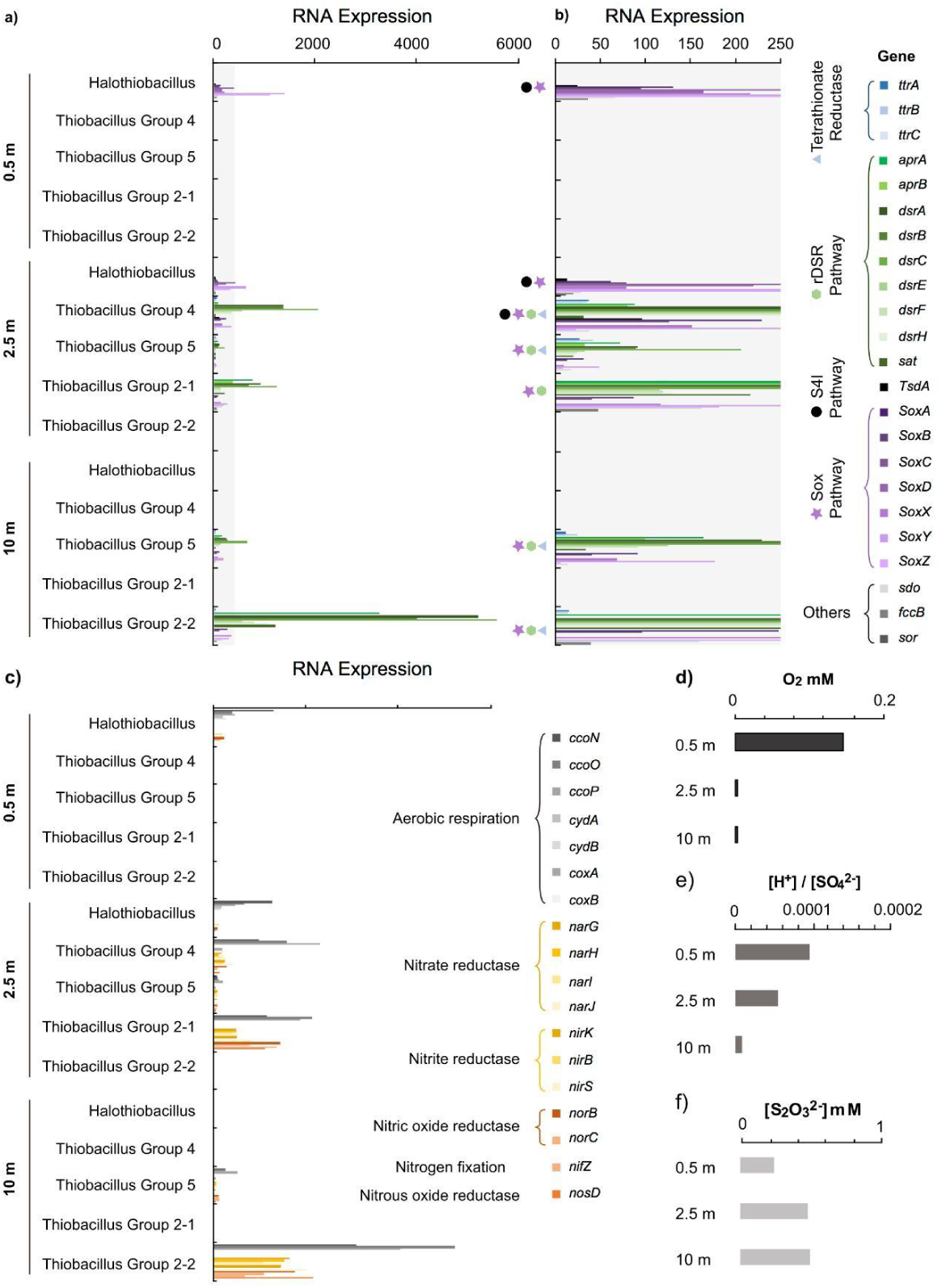
RNA-seq analysis of gene expression of *Halothiobacillus* and *Thiobacillus* (groups 4, 5, 2-1, 2-2). Samples collected from August 2018 at the depths of 0.5 m (epilimnion), 2.5 m (upper metalimnion) and 10 m (hypolimnion) were analyzed. Gene expression levels of (a) tetrathionate reductase genes (*ttrABC*), rDSR pathway genes (*aprAB, dsrABCEFH, sat*), S4I pathway genes (*tsdA*), Sox pathway genes (s*oxABCDXYZ*) and *sdo, fccB* and *sor*, (b) (zoom in at the 0 – 250 gene expression level of (a) to show tetrathionate reductase (*ttrABC*) and S4I pathway (*tsdA*) depth dependent observed activities, and (c) aerobic respiration (*ccoNOP, cydAB*, c*oxAB*) and nitrogen reductive genes (*narGHIJ, nirKBS, norBC* and *nosS*). The (d) O_2_ (mM) concentrations, (e) H^+^/SO_4_^2-^ ratios and (f) S_2_O_3_^2-^ (mM) concentrations at the three depths.

This depth dependent segregation of sulfur oxidizing genera and thus sulfur pathways associated with decreasing oxygen concentrations (Figure 7d), correlates with a clear differentiation of sulfur cycling outcomes associated with the shift from *Halothiobacillus* in the upper more oxygenated waters to *Thiobacillus* in the metalimnetic micro-oxic (Groups 1 and 5) and deeper anoxic waters (Group 2). Specifically, decreasing acidity generation (Figure 7e) and increasing thiosulfate concentrations (Figure 7f) occurred with depth.

## Discussion

### Neutrophilic chemolithoautotrophic sulfur oxidizing bacteria dominate tailing impoundments waters over time and space

The results demonstrate clearly the importance of neutrophilic chemolithoautotrophic sulfur oxidizing bacteria in driving sulfur compound oxidation and acidification within tailing impoundment waters. The tailing impoundment water was consistently inhabited by neutrophilic chemolithoautotrophic sulfur oxidizing bacteria and (photo)-heterotrophs (Figure 3a). The sulfur oxidizing portion of the communities included members of the genera *Halothiobacillus, Sulfuricurvum, Thiobacillus, Thiovirga, Sulfurovum* and *Brevundimonas* (Figure 3a) that have been previously implicated in sulfur oxidation ^15,21,67,70–85^. Members of the *Sediminibacterium* genus have been reported as iron oxidizing bacteria ^76,86,87^; however, here metagenomic analyses found that they possess the *sdo* gene, suggesting they could be involved in sulfur oxidation as well.

Notably, the microbial community inhabiting the tailing impoundment waters (pH ∼ 7 to 4) shared few similarities with more well studied mining associated ARD communities present under more acidic conditions (e.g.,^11,88^). This result is not surprising given that pH has a strong effect on microbial community structure ^33^. None of the typical ferrous and sulfur oxidizing species found to dominate in acidic mining waters (e.g., *Acidithiobacillus thiooxidans, Acidithiobacillus ferrooxidans* ^31^) occurred in these circumneutral high sulfur waters (Figure 3a). However, the communities analyzed in this study possessed similar genera as systems experiencing accelerating metal/concrete corrosion^29^. Further, Lopes et al.^32^ identified both *Halothiobacillus* and *Thiobacillus* as dominant genera present in circumneutral processing waters at active mine sites in Portugal, suggesting that these sulfur oxidizers are highly adapted to these high metal and sulfur environments globally. With the exception of certain members of *Thiobacillus,* such as *Thiobacillus ferrooxidans,* whose presence and activity has been noted in various acid mine drainage systems ^89^, the dominant neutrophilic sulfur oxidizing genera (*Halothiobacillus, Sulfuricurvum, Sediminibacterium* and other *Thiobacillus* spp.) identified in this study have largely gone underappreciated as important organisms mediating oxygen consumption, sulfur compound transformation and acidity generation in mining impacted waters. In particular, the relationships between their distributions, metabolic functional genes and the geochemical outcomes in environmental settings have not been well delineated underlining the novelty of our work.

### *Halothiobacillus* is an important driver of acidity generation via direct aerobic sulfur oxidation

The relative proportions of various sulfur oxidizing bacteria and community level occurrence of sulfur metabolic genes correlated strongly with different sulfur geochemical outcomes, particularly with generation of acidity and thiosulfate, which is a key tailings associated sulfur compound and important sulfur substrate for sulfur oxidizing genera. Coincident to the years when *Halothiobacillus* was the dominant sulfur oxidizing bacteriium (2016 and 2017), lower thiosulfate concentrations and significantly higher H^+^/SO_4_^2-^ ratios occurred in the water column (Figures 1-3). The *Halothiobacillus* genus likely directly oxidizes reduced sulfur compounds to sulfate (Figure 6) without formation of free sulfur oxidation intermediates through the Sox pathway ^16,90^. However, as *Halothiobacillus* also encodes the *tsdA* gene (Figures 5 and 6), this organism may also use the S4I pathway, and this is supported by expression of the *Halothiobacillus* t*sdA* gene in the epilimentic (oxygenated) and metalimnetic (low O_2_) region of the water cap (Figure 7). *Halothiobacillus* also has been detected in moderately acidic (pH∼4) waters discharged from an active mine into the receiving environment ^15^.

Our *Halothiobacillus*-dominated enrichments rapidly drove circumneutral pH conditions to acidic levels under aerobic conditions ^15,91^. Previous studies have shown that acidity may be generated by the production of CO_2_ during the respiration of organic matter by heterotrophic communities ^92^ and slightly lower organic carbon concentration were observed in 2016 and 2017 (Table S3) thus some acidity generation may be attributed to that. However, organic carbon concentrations are relatively low in this system compared to reduced sulfur. In addition, the enrichment and genomic results provide strong evidence that *Halothiobacillus* is capable of driving circumneutral conditions to lower pH through a direct sulfur oxidation pathway, resulting in sulfate and acid in mine water systems when oxygen is present. *Sulfuricurvum* found in this system, like *Halothiobacillus*, also contains genes for the complete Sox complex (s*oxABCDXYZ*) and no rDSR or HDR encoding genes (Figure 5) and its relative abundance in 2015 was similar to that of *Halothiobacillus* in 2016/2017 (Figure 1). However, its presence in 2015 did not coincide with any significant acidity generation (Figure 1 and 2).

In 2016 and 2017, more oxygenated conditions may have enabled *Halothiobacillus* to outcompete other members of the sulfur oxidizing community such as *Thiobacillus* and *Sediminibacterium* (Figure 3a). *Halothiobacillus* has been proposed as an early indicator of net acid generating conditions in circumneutral waters ^15,91^. This study strongly supports the notion as higher spring oxygen concentrations and proliferation of *Halothiobacillus* preceded a progressive increase in acid production on the order of months in 2016 and 2017. However, the relationship between higher spring oxygen concentrations and the presence of *Halothiobacillus* does not explain its continued dominance as anoxic/micro-oxic conditions are established in the metalimnion and hypolimnion during summer stratification (Figure 3). Emerging evidence has shown that obligate aerobe*s* under anoxic conditions may participate in syntrophic cryptic oxygen cycling utilizing a heterotrophic partner that can generate molecular oxygen (i.e. through perchlorate respiration) ^93,94^. However, to the best of our knowledge, the family Sphingomonadaceae does not have demonstrated perchlorate respiring abilities nor was perchlorate measured directly in this study ^95^. In addition, RNA-seq analysis suggested that *Halothiobacillus* and members of the Sphinogmonadaceae family were not metabolically active in August 2018 at the time of RNA sampling in the depleted oxygen zone, but *Thiobacillus* was shown to express aerobic respiration genes (Figure 7). Thus, the metabolic activity of obligate aerobes where no oxygen or little oxygen is detectable in tailing impoundments remains speculative and warrants more research.

### Niche partitioning of Sox and rDSR bacterial oxidation strategies across steep oxygen gradients

The high amounts of reduced sulfur inputs into this system (sulfide tailing deposition and processing waters) resulted in significant oxygen consumption and steep gradients throughout the water column (Figure 2b). These provide an opportunity to examine ecological niche partitioning of differing bacterial metabolic strategies to carry out sulfur oxidation through various pathways (i.e. Sox vs. rDSR) and the capacity of those bacteria to couple oxidation to terminal electron acceptors like the O_2_, NO_3_^-^ and NO_2_^-^. Consistent with the connection with oxygenated conditions observed over time in this study, RNA-seq analysis in August 2018 confirmed that *Halothiobacillus* metabolic activity was limited to the epilimnetic and to a lesser degree, the upper metalimnetic portion of the water column, where higher oxygen concentrations were present (Figure 7). Conversely, in 2015 and 2018, *dsrEBFH, aprAB, sat* and genes encoding incomplete Sox complex (lacking SoxCD) (possessed by *Thiobacillus*) were progressively more expressed correlating with depth and the steep decreasing oxygen gradient that begins in the metalimnetic region of the tailings impoundment (Figure 4a). Phylogenetic analysis comparing DsrAB proteins in our ecosystem to organisms shown to utilize the rDSR pathway suggested that these proteins are being utilized for oxidative reactions by *Thiobacillus* rather than for reduction (Figure S1). Therefore, our results are well aligned with recent work showing that environmentally relevant chemolithoautotrophic sulfur cycling bacteria can mediate a reversal of the sulfate reduction pathway catalyzed through the Dsr complex to disproportionate, or oxidize reduced sulfur species ^79,96,97^. This capability has been also identified in phototrophic sulfur bacteria such as the *Allochromatium vinosum* (*Chromatiaceae* family)^97 96^.

Metagenomic characterization of sulfur oxidizing bacteria from this tailings impoundment strongly supports that rDSR pathways are much slower in generating acidity and sulfate compared to the Sox pathway, and there is potential for free sulfite formation through rDSR (Figure 4b). *In situ* analysis of sulfite concentrations in the tailings impoundment water did not provide evidence consistent with increased sulfite formation when rDSR pathways were more abundant (Table S4), but sources of sulfite beyond the rDSR pathways (i.e. directly from mill effluents) can overprint sulfite formation and utilization from biogeochemical cycling, particularly at these relatively low concentrations (Table S2). In addition, the rDSR pathways coincided with the presence of genes encoding TtrABC (Figure 4a), which may reduce tetrathionate, regenerating thiosulfate, as well as with an incomplete Sox pathway (lack of SoxCD). The rDSR pathways are proposed to be more energy efficient than Sox pathways^28^ likely evolving in the early Archean era when the ability to oxidize sulfur compounds under anoxic conditions would have been necessary ^25^. Their increased prevalence with depth (Figure 4a;S3) and higher gene expression levels (Figure 7) are consistent with these pathways dominating in the micro/anoxic depths of these tailing impoundment waters and their reactions may be coupled to both aerobic respiration and nitrate/nitrous oxide reduction (Figure 7). *Thiobacillus* expressed nitrate reduction and aerobic respiration genes at increased depth in August 2018. However, ammonia concentrations did not increase with depth (Table S6), likely reflecting multiple sources of ammonia, such as ammonia nitrate from rock blasting or a by-product of the cyanidation process during ore recovery in mill processing. These allochthonous N sources are contributed during on-going discharges of mine impacted waters and tailings to the system, which can overprint a possible signal from denitrification. Nevertheless, the RNA-seq results (Figure 7) suggest that *s*everal relatively abundant populations of *Sulfuricurvum* and *Thiobacillus* observed during 2015 and 2018 summer months (Figure 3) may be coupling sulfur oxidation to denitrification under lower oxygen concentrations occurring deeper in the impoundment water column across time in this system. The reactions catalyzed by these different pathways (Figure 5) will determine the distribution and transformation of sulfur species within the impoundment, but would be cryptic from a sulfate generation standpoint. These biological sulfur oxidation pathways are currently not accounted for in model predictions but are likely important in determining when and where higher concentrations of reduced sulfur compounds such as thiosulfate will occur. As tailing impoundments are commonly a treatment stage prior to mining water being discharged into the environment, accumulations of reduced phases of sulfur in discharged waters can impact receiving environments through subsequent acidity generation and oxygen consumption.

An adaptive capability of indigenous organisms to couple sulfur oxidation to nitrate or nitrous oxide reduction is likely an energetically favourable and viable metabolic strategy in tailing impoundment waters where sulfur oxidation has driven waters micro-oxic or anoxic (albeit with simultaneous potential for cryptic aerobic respiration discussed above) and oxidized nitrogen species are present (Table S6). Experimentally it is well established that *Thiobacillus denitrificans* can utilize nitrate as a terminal electron acceptor^78^. In other studies, *Desulfurivibrio alkaliphilus* was shown to be capable of oxidizing reduced sulfur species coupled to nitrate reduction using rDSR under anoxic conditions ^96^. Similarly, in the deep terrestrial subsurface, the coupling of nitrate reduction and S oxidation via rDSR mediation has also been found to occur ^98,99^. Thus, while there was a consistent presence of *soxABCXZ* and *tsdA* (*Halothiobacillus, Sulfuricurvum*) throughout depth and across time in this study, it appears that *Thiobacillus* may gain a competitive advantage over *Halothiobacillus* under low oxygen to anoxic conditions, using the more energy efficient rDSR pathways coupled to denitrification or small amounts of oxygen when available.

## Conclusion

This study demonstrates that oxygen driven niche partitioning of microbial sulfur oxidation pathways results in environmentally important outcomes such as acidity generation and thiosulfate persistence in circumneutral mining waters. In the absence of detectable oxygen or low oxygen conditions, chemolithoautotrophic sulfur oxidizers, such as members of the *Thiobacillus* genus, are highly active via sulfur oxidation coupled to nitrate/O_2_ through the efficient rDSR pathway. Conversely, *Halothiobacillus* prevailed under oxygenated conditions and its proliferation preceded significant acidity generation on the order of months. Clear linkages were demonstrated between oxygen status, sulfur oxidizing genera, sulfur pathways, and resulting sulfur biogeochemical cycling outcomes in the mine tailings impoundment waters. Further, SOB enrichment experiments for both this impoundment as well as three other mine impoundment waters previously showed the same patterns indicating the wider relevance of our findings. These results identify that the analysis of microbial genes and/or of microbial community distributions is a promising avenue to pursue for accurate prediction of whether a system is moving towards net acidity generation or accumulating higher concentrations of sulfur oxidation intermediates. In addition, there may be an opportunity for these organisms to be used within *in situ* bioengineered management or treatment strategies (e.g., by amplifying or decreasing sulfur oxidative pathways). The results of this study reveal SOB occurring in actively managed mine tailings impoundments, extend our understanding of ecological partitioning of sulfur biogeochemical cycling and illuminate the possibility of biological strategies to mitigate mining impacted water challenges.

## Acknowledgements

The authors thank all of the on-site mine personnel who aided in site orientation, sample collection and processing. Research was supported by the Genome Canada Large Scale Applied Research Program and Ontario Research Fund–Research Excellence grants to LW and JB. The National Sciences and Engineering Research Council CREATE Mine of Knowledge post-doctoral fellowship to KW-M is also gratefully acknowledged.

