## Supplementary Material for "Acidity and sulfur oxidation intermediate concentrations controlled by O_2_-driven partitioning of sulfur oxidizing bacteria in a mine tailings impoundment"

**Table S1.** Physiochemical conditions across time (2015-2018) (21 sampling campaigns, n=53) of the Northern Ontario tailings impoundment water (temperature (°C), pH, conductivity (µS/cm$^2$), oxidation-reduction potential (ORP, mV), $O_2$ % and $O_2$ (mg/L). NA indicates that data is not available.

| Date | Depth (m) | °C | pH | Conductivity (µS/cm$^2$) | ORP | $O_2$% | $O_2$ (mg/L) |
|---|---|---|---|---|---|---|---|
| March 2015 | 0 | 0.4 | 8.32 | 2073 | 84 | 13.3 | 1.91 |
| | 0.75 | 3.3 | 8.51 | 2238 | 63 | 0.8 | 0.1 |
| | 21 | 3.2 | 8.76 | 2241 | 32 | 0.2 | 0.03 |
| May 2015 | 2.5 | 8.9 | 7.05 | 2130 | 150 | 21 | 2.41 |
| | 3.5 | 8.5 | 7.01 | 2135 | 141 | 10 | 1.16 |
| | 15 | 7.3 | 7.21 | 2147 | 138 | 2.5 | 0.3 |
| July 10 2015 | 2 | 15.4 | 7.28 | 1803 | 63.1 | 12.5 | 1.23 |
| | 4 | 14.5 | 8.25 | 1757 | -94 | 3.9 | 0.39 |
| | 30 | 10.5 | 8.45 | 1581 | -289 | 0.4 | 0.04 |
| July 22 2015 | 1 | 19.4 | 6.97 | 2246 | 187 | 63.1 | 5.77 |
| | 3.5 | 17.5 | 7.28 | 2235 | 177 | 10.8 | 1 |
| | 10 | 14.3 | 8.34 | 2223 | 133 | 2.5 | 0.26 |
| August 2015 | 1 | 18.5 | 6.85 | 2069 | 203 | 54.2 | 5.04 |
| | 2.8 | 16.8 | 7 | 2069 | 203 | 0.6 | 0.05 |
| | 26 | 14.8 | 7.35 | 2056 | 193 | <LOD | <LOD |
| September 2015 | 1 | 16.5 | 6.8 | 2198 | 159 | 19.1 | 1.85 |
| | 7 | 16.5 | 6.68 | 2199 | 129 | 16.3 | 1.58 |
| | 33 | 13 | 6.73 | 2162 | 43 | 3.7 | 0.37 |
| November 2015 | 1.3 | 11 | 6.35 | 2220 | 216 | 39.4 | 4.31 |
| May 2016 | 1 | 9.0 | 6.4 | 1909 | 152 | 55 | 6.3 |
| | 6 | 6.6 | 6.46 | 1919 | 170 | 33.5 | 4.09 |
| July 2016 | 0.5 | 22.3 | 4.79 | 1990 | 250 | 74.5 | 6.44 |
| | 1.5 | 17 | 5.83 | 2013 | 205 | 5.9 | 0.57 |
| | 10 | 15.8 | 6.28 | 2009 | 148 | 0.5 | 0.06 |
| September 2016 | 1 | 19 | 4.69 | 2097 | 239 | 43.6 | 4.04 |
| | 6 | 18.9 | 4.75 | 2096 | 243 | 27.9 | 2.58 |
| | 32 | 15.3 | 6.25 | 2067 | -46 | 21.8 | 2.19 |
| May 2017 | 0.5 | 10.7 | NA | 2109 | 70 | 31.6 | 3.49 |
| June 2017 | 0.5 | 16.8 | 5.49 | 2098 | 181 | 57.9 | 5.39 |
| | 2 | 15.6 | 5.76 | 2125 | 177 | 32 | 3.49 |

| | | | | | | | |
|---|---|---|---|---|---|---|---|
| | 10 | 12 | 6.63 | 2145 | 158 | 0.6 | 0.06 |
| July 2017 | 0.5 | 19.1 | 5.43 | 2130 | 214 | 61.4 | 5.16 |
| | 2 | 15.3 | 5.81 | 2163 | 192 | 24.4 | 1.72 |
| | 10 | 14.1 | 6.43 | 2162 | 152 | 4.2 | 0.43 |
| August 2017 | 0.5 | 19.2 | 5.26 | 2186 | 207 | 61.4 | 10.26 |
| | 2 | 17.6 | 5.09 | 2245 | 204 | 24.4 | 2.78 |
| | 10 | 15.7 | 6.78 | 2194 | 158 | 4.2 | 0.29 |
| September 2017 | 0.5 | 21.4 | 4.67 | 2288 | 212 | 72.6 | 6.38 |
| | 2 | 20.7 | 4.73 | 2299 | 219 | 45.4 | 4.16 |
| | 10 | 17 | 6.46 | 2287 | 76 | <LOD | <LOD |
| November 2017 | 0.5 | 11.3 | 6.94 | 2097 | 111 | 30.6 | 2.76 |
| | 10 | 11.3 | 6.15 | 2091 | 117 | 22 | 2.33 |
| May 2018 | 0.5 | 12 | 6.94 | NA | 159 | 57.2 | 6.12 |
| | 2.8 | 6.1 | 7.66 | NA | 149 | 11.4 | 1.39 |
| | 21 | 4.5 | 8.27 | NA | 168 | 0.3 | 0.04 |
| June 2018 | 0.5 | 12.3 | 6.75 | 2227 | 187 | 12.8 | 1.29 |
| July 11 2018 | 0.5 | 21.4 | 6.86 | 2221 | 180 | 65.8 | 5.8 |
| | 2.8 | 20.7 | 6.74 | 2218 | 178 | 62.4 | 5.56 |
| | 10 | 12.8 | 7.2 | 2193 | 163 | 1.4 | 0.14 |
| July 23 2018 | 0.5 | 21.7 | 8.18 | 2175 | 199 | 57.8 | 5.05 |
| August 2018 | 0.5 | 22.5 | 6.08 | 2227 | 163 | 54.2 | 4.68 |
| | 2.5 | 18.2 | 6.33 | 2187 | 135 | 0.6 | 0.05 |
| | 10 | 16.8 | 7.16 | 2199 | 152 | 0.2 | 0.02 |

**Table S2.** Sulfur speciation (n=53) (mmol/L) across time (2015-2018) of the tailings impoundment waters ($TotS_{0.2µm}$, $TotS_{0.45µm}$, $TotS_{UF}$, $S\text{-}SO_4^{2-}$, $S\text{-}SO_3^{2-}$, $S\text{-}S_2O_3^{2-}$, $S\text{-}S_3O_6^{2-}$ (where available), $S\text{-}S_4O_6^{2-}$ (where available) $ΣH_2S$ and Sreact (defined as $TotS_{0.45µm}$ - $S\text{-}SO_4^{2-}$).

| Date | Depth (m) | $TotS_{0.2 µm}$ mmol/L | $TotS_{0.45 µm}$ mmol/L | $TotS_{UF}$ mmol/L | $S\text{-}SO_4^{2-}$ mmol/L | $S\text{-}SO_3^{2-}$ mmol/L | $S\text{-}S_2O_3^{2-}$ mmol/L | $S\text{-}S_3O_6^{2-}$ mmol/L | $S\text{-}S_4O_6^{2-}$ mmol/L | $S\text{-}ΣH_2S$ µmol/L | $S_{react}$ mmol/L |
|---|---|---|---|---|---|---|---|---|---|---|---|
| March 2015 | 0 | 8.0 ± 0.04 | 8.0 ± 0.7 | - | 6.5 ± 0.3 | 0.24 ± 0.5 | 0.26 ± 0.16 | - | - | <0.01 | 1.5 |
| | 0.75 | 9.7 ± 0.08 | 8.8 ± 0.03 | - | 6.6 ± 0.3 | <LOD | 0.12 ± 0.01 | - | - | <0.01 | 2.2 |
| | 21 | 9.8 ± 0.05 | 9.9 ± 0.05 | - | 6.6 ± 0.3 | 0.05 ± 0.05 | 0.59 ± 0.41 | - | - | 1 ± 0.02 | 3.3 |
| May 2015 | 2.5 | 8.3 ± 0.04 | 8.3 ± 0.02 | - | 8.5 ± 0.5 | <LOD | 0.26 ± 0.05 | - | - | 0.14 ± 0.01 | <LOD |
| | 3.5 | 8.4 ± 0.03 | 8.3 ± 0.02 | - | 8.5 ± 0.5 | <LOD | 0.18 ± 0.04 | - | - | 0.29 ± 0.02 | <LOD |
| | 15 | 8.4 ± 0.03 | 8.4 ± 0.03 | - | 8.6 ± 0.5 | <LOD | 0.16 ± 0.02 | - | - | 0.44 ± 0.02 | <LOD |
| July 10 2015 | 2 | 8.2 ± 0.01 | 8.2 ± 0.02 | - | 2.6 ± 0.5 | <LOD | 0.20 ± 0.04 | - | - | 0.24 ± 0.01 | 5.6 |
| | 4 | 8.4 ± 0.04 | 8.4 ± 0.01 | - | 1.5 ± 0.5 | <LOD | 0.23 ± 0.04 | - | - | 0.65 ± 0.06 | 6.9 |
| | 30 | 8.3 ± 0.03 | 8.3 ± 0.01 | - | 2 ± 0.5 | <LOD | 0.14 ± 0.001 | - | - | 0.75 ± 0.04 | 6.3 |
| July 22 2015 | 1 | 8.3 ± 0.01 | 8.3 ± 0.01 | - | 1.4 ± 0.5 | <LOD | 0.10 ± 0.35 | - | - | 0.19 ± 0.01 | 6.9 |
| | 3.5 | 8.4 ± 0.01 | 8.4 ± 0.01 | - | 2.2 ± 0.5 | <LOD | 0.10 ± 0.02 | - | - | 0.18 ± 0.01 | 6.2 |
| | 10 | 8.5 ± 0.05 | 8.4 ± 0.01 | - | 1.7 ± 0.5 | <LOD | 0.10 ± 0.04 | - | - | 0.37 ± 0.03 | 6.7 |
| August 2015 | 1 | 8.3 ± 0.02 | 8.3 ± 0.02 | - | 8.4 ± 0.5 | 0.14 ± 0.02 | 0.27 ± 0.01 | - | - | < 0.01 | <LOD |
| | 2.8 | 8.5 ± 0.04 | 8.5 ± 0.02 | - | 9.1 ± 0.5 | 0.16 ± 0.004 | 0.22 ± 0.02 | - | - | <0.01 | <LOD |
| | 26 | 8.5 ± 0.01 | 8.5 ± 0.01 | - | 7.5 ± 0.5 | 0.15 ± 0.004 | 0.21 ± 0.01 | - | - | 0.73 ± 0.05 | 1 |
| Sept 2015 | 1 | 9.0 ± 0.1 | 9.1 ± 0.04 | - | 6.8 ± 0.5 | <LOD | 0.04 ± 0.01 | - | - | <0.01 | 2.3 |
| | 7 | 9.0 ± 0.03 | 8.8 ± 0.01 | - | 6.9 ± 0.5 | <LOD | 0.03 ± 0.001 | - | - | <0.01 | 1.9 |
| | 33 | 8.7 ± 0.02 | 8.8 ± 0.03 | - | 6.8 ± 0.5 | <LOD | 0.02 ± 0.02 | - | - | 1.5 ± 0.02 | 2 |
| Nov | 1.3 | 8.7 ± | 8.8 ± 0.07 | - | 6.8 ± 0.5 | 0.04 ± | 0.26 ± | - | - | 0.2 ± 0.02 | 2 |

| | | | | | | | | | | | |
|---|---|---|---|---|---|---|---|---|---|---|---|
| 2015 | | 0.02 | | | | 0.003 | 0.02 | | | | |
| May 2016 | 1 | 7.8 ± 0.13 | 7.6 ± 0.04 | - | 5.4 ± 0.5 | 0.04 ± 0.004 | 0.12 ± 0.07 | - | - | 1.3 ± 0.05 | 2.2 |
| | 6 | 7.9 ± 0.08 | 7.9 ± 0.01 | - | 5.7 ± 0.5 | 0.03 ± 0.006 | 0.12 ± 0.07 | - | - | 1.2 ± 0.05 | 2.2 |
| July 2016 | 0.5 | 8.2 ± 0.07 | 8.0 ± 0.17 | 8.1 ± 0.05 | 6.1 ± 0.5 | 0.024 ± 0.004 | 0.06 ± 0.01 | - | - | <0.01 | 1.9 |
| | 1.5 | 8.3 ± 0.06 | 8.4 ± 0.18 | 8.3 ± 0.13 | 6.1 ± 0.5 | 0.013 ± 0.006 | 0.08 ± 0.01 | - | - | 0.59 ± 0.02 | 2.3 |
| | 10 | 8.3 ± 0.1 | 8.5 ± 0.07 | 8.4 ± 0.08 | 6.2 ± 0.5 | 0.012 ± 0.004 | 0.14 ± 0.03 | - | - | <0.01 | 2.3 |
| Sept 2016 | 1 | 8.7 ± 0.12 | 8.9 ± 0.08 | 8.8 ± 0.03 | 6.2 ± 0.5 | 0.04 ± 0.004 | 0.12 ± 0.03 | - | - | 0.44 ± 0.06 | 2.7 |
| | 6 | 8.7 ± 0.15 | 8.8 ± 0.09 | 8.7 ± 0.15 | 6.6 ± 0.5 | 0.03 ± 0.01 | 0.09 ± 0.03 | - | - | 0.44 ± 0.03 | 2.2 |
| | 32 | 8.7 ± 0.06 | 8.6 ± 0.22 | 8.5 ± 0.12 | 6.5 ± 0.5 | 0.05 ± 0.01 | 0.14 ± 0.01 | - | - | 6 ± 0.4 | 2.1 |
| May 2017 | 0.5 | 8.8 ± 0.06 | 8.8 ± 0.03 | 8.9 ± 0.09 | 1.8 ± 0.6 | 0.04 ± 0.01 | 0.05 ± 0.01 | - | - | <0.01 | 7 |
| June 2017 | 0.5 | 8.7 ± 0.12 | 8.6 ± 0.1 | 8.6 ± 0.15 | 7.0 ± 0.5 | 0.02 ± 0.01 | 0.087 ± 0.02 | - | - | <0.01 | 1.6 |
| | 2 | 8.8 ± 0.17 | 8.7 ± 0.1 | 8.7 ± 0.06 | 7.0 ± 0.5 | 0.02 ± 0.002 | 0.025 ± 0.03 | - | - | <0.01 | 1.7 |
| | 10 | 9.1 ± 0.16 | 9.1 ± 0.14 | 9.3 ± 0.14 | 8.2 ± 0.5 | 0.04 ± 0.02 | 0.16 ± 0.06 | - | - | 0.83 ± 0.05 | 0.9 |
| July 2017 | 0.5 | 8.9 ± 0.11 | 9.1 ± 0.07 | 8.9 ± 0.08 | 6.9 ± 0.5 | 0.05 ± 0.02 | 0.11 ± 0.05 | - | - | <0.01 | 2.2 |
| | 2 | 9.2 ± 0.06 | 9.3 ± 0.13 | 9.3 ± 0.08 | 7.1 ± 0.5 | 0.03 ± 0.01 | 0.13 ± 0.01 | - | - | <0.01 | 2.2 |
| | 10 | 9.3 ± 0.04 | 9.5 ± 0.08 | 9.3 ± 0.14 | 7.3 ± 0.5 | 0.03 ± 0.01 | 0.24 ± 0.02 | - | - | 0.64 ± 0.05 | 2.2 |
| August 2017 | 0.5 | 8.7 ± 0.06 | 8.8 ± 0.14 | 8.6 ± 0.03 | 9.8 ± 1.3 | 0.03 ± 0.002 | 0.09 ± 0.01 | - | - | - | <LOD |
| | 2 | 9.0 ± 0.34 | 9.3 ± 0.13 | 9.3 ± 0.07 | 10.2 ± 0.5 | 0.02 ± 0.002 | 0.02 ± 0.01 | - | - | - | <LOD |
| | 10 | 9.4 ± 0.14 | 9.3 ± 0.08 | 9.2 ± 0.11 | 10.2 ± 0.5 | 0.03 ± 0.005 | 0.11 ± 0.001 | - | - | - | <LOD |
| Sept 2017 | 0.5 | 10.5 ± 0.26 | 10.4 ± 0.15 | 10.7 ± 0.22 | 8.9 ± 0.5 | 0.007 ± 0.001 | 0.01 ± 0.001 | - | - | 0.94 ± 0.05 | 1.5 |
| | 2 | 10.8 ± 0.5 | 10.5 ± 0.16 | 10.8 ± 0.08 | 7.0 ± 0.5 | 0.008 ± 0.001 | 0.01 ± 0.001 | - | - | 1.0 ± 0.08 | 3.5 |
| | 10 | 10.1 ± 0.7 | 9.6 ± 0.03 | 9.6 ± 0.04 | 7.3 ± 0.5 | 0.008 ± 0.001 | 0.04 ± 0.001 | - | - | 1.5 ± 0.06 | 2.3 |
| Nov 2017 | 0.5 | 9.4 ± 0.1 | 9.4 ± 0.11 | 9.5 ± 0.10 | 8.4 ± 0.5 | 0.011 ± 0.001 | 0.03 ± 0.001 | - | - | 0.5 ± 0.03 | 1 |

| | | | | | | | | | | | |
|---|---|---|---|---|---|---|---|---|---|---|---|
| | 10 | 9.5 ± 0.1 | 9.5 ± 0.08 | 9.4 ± 0.11 | 0.27 ± 0.5 | 0.009 ± 0.001 | 0.02 ± 0.01 | - | - | 0.6 ± 0.04 | 9.2 |
| May 2018 | 0.5 | - | - | - | 6.1 ± 0.5 | - | - | 0.08 | 0.1 | - | - |
| | 2.8 | - | 7.7 ± 0.06 | - | 7.6 ± 0.5 | 0.009 ± 0.002 | 0.08 ± 0.003 | 0.09 | 0.1 | - | 0.1 |
| | 21 | 9.1 ± 0.2 | 9.2 ± 0.12 | 9.3 ± 0.06 | - | 0.012 ± 0.002 | 0.29 ± 0.04 | <0.03 | 0.01 | - | - |
| June 2018 | 0.5 | - | 9.5 ± 0.04 | - | 7.7 ± 0.5 | 0.015 ± 0.008 | 1.7 ± 0.7 | 0.03 | 0.03 | 1.2 ± 0.3 | 2.1 |
| July 11 2018 | 0.5 | - | 9.7 ± 0.07 | - | 8.1 ± 0.5 | 0.007 ± 0.002 | 0.13 ± 0.01 | - | - | <0.01 | 1.6 |
| | 2.8 | | - | - | | - | - | <0.03 | 0.09 | 0.67 ± 0.08 | 1.05 |
| | 10 | - | 9.9 ± 0.03 | - | 10.2 ± 2.2 | 0.017 ± 0.001 | 0.38 ± 0.03 | - | - | 1.7 ± 0.2 | <LOD |
| July 23 2018 | 0.5 | - | 9.6 ± 0.02 | - | 7.9 ± 0.7 | - | - | - | - | <0.01 | 2.1 |
| August 2018 | 0.5 | - | 9.8 ± 0.07 | - | 8.6 ± 0.5 | 0.007 ± 0.003 | 0.23 ± 0.21 | <0.03 | 0.1 | 0.58 ± 0.1 | 1.2 |
| | 2.5 | - | 9.8 ± 0.05 | - | 8.8 ± 0.5 | 0.001 ± 0.003 | 0.48 ± 0.05 | <0.03 | 0.02 | 1.2 ± 0.1 | 1 |
| | 10 | 9.9 ± 0.1 | 10.1 ± 0.14 | 0.2 ± 0.07 | 8.9 ± 0.5 | 0.004 ± 0.004 | 0.50 ± 0.05 | <0.03 | 0.02 | 17 ± 0.2 | 1.2 |

**Table S3**. 16SrRNA amplicons for mining waters (n=30) with Shannon's Diversity Index, Pielou's Eveness (J), richness and total sequences.

| | Depth (m) | Year | Month | Shannon Diversity (H') | Pielou's Evenness (J) | Richness | Total Sequences |
|---|---|---|---|---|---|---|---|
| March 2015 0 m | 0 | 2015 | March | 2.5 | 0.22 | 120 | 101253 |
| March 2015 0.75 m | 0.75 | 2015 | March | 2.6 | 0.22 | 77 | 90750 |
| March 2015 21 m | 21 | 2015 | March | 2.51 | 0.22 | 71 | 87785 |
| May 2015 2.5 m | 2.5 | 2015 | May | 1.9 | 0.17 | 119 | 86194 |
| May 2015 3.5 m | 3.5 | 2015 | May | 1.79 | 0.16 | 83 | 61878 |
| August 2015 1 m | 1 | 2015 | August | 3.3 | 0.28 | 357 | 141767 |
| August 2015 2.8 m | 2.8 | 2015 | August | 2.99 | 0.27 | 155 | 58777 |
| August 2015 25.8 m | 25.8 | 2015 | August | 2.31 | 0.2 | 126 | 94984 |
| July 2016 1.5 m | 1.5 | 2016 | July | 2.01 | 0.17 | 142 | 132304 |
| July 2016 10 m | 10 | 2016 | July | 1.85 | 0.16 | 111 | 113926 |
| June 2017 2 m | 2 | 2017 | June | 2.23 | 0.19 | 210 | 128340 |
| June 2017 10 m | 10 | 2017 | June | 1.63 | 0.14 | 72 | 124093 |
| July 2017 0.5 m | 0.5 | 2017 | July | 2.8 | 0.24 | 327 | 123386 |
| July 2017 2 m | 2 | 2017 | July | 2.27 | 0.2 | 221 | 98468 |
| July 2017 10 m | 10 | 2017 | July | 1.49 | 0.13 | 78 | 110501 |
| August 2017 0.5 m | 0.5 | 2017 | August | 2.4 | 0.2 | 392 | 131216 |
| August 2017 2 m | 2 | 2017 | August | 1.88 | 0.16 | 305 | 127625 |
| August 2017 10 m | 10 | 2017 | August | 1.51 | 0.13 | 92 | 112451 |
| September 2017 10 m | 10 | 2017 | September | 1.36 | 0.12 | 97 | 108581 |
| November 2017 0.5 m | 0.5 | 2017 | November | 1.94 | 0.17 | 149 | 108630 |
| November 2017 10 m | 10 | 2017 | November | 1.94 | 0.16 | 184 | 129863 |
| May 2018 0.5 m | 0.5 | 2018 | May | 3.4 | 0.32 | 401 | 52146 |
| May 2018 2.8 m | 2.8 | 2018 | May | 3.7 | 0.36 | 103 | 28650 |
| May 2018 10 m | 10 | 2018 | May | 1.65 | 0.15 | 71 | 50707 |
| July 2018 0.5 m | 0.5 | 2018 | July | 2.7 | 0.22 | 548 | 240164 |
| July 2018 10 m | 10 | 2018 | July | 2.73 | 0.23 | 235 | 137778 |
| July 23 2018 0.5 m | 0.5 | 2018 | July | 4.9 | 0.4 | 1151 | 142624 |
| August 2018 0.5 m | 0.5 | 2018 | August | 2.6 | 0.21 | 454 | 202000 |
| August 2018 2.5 m | 2.5 | 2018 | August | 2.95 | 0.24 | 209 | 195046 |
| August 2018 10 m | 10 | 2018 | August | 2.27 | 0.18 | 176 | 211143 |

**Table S4.** Mine tailing impoundment aquatic microbial community composition within three statistical clusters 1) 2015 (n=8 2) 2016/2017 (n= 3 and 13) 2018 (n= 9) as determined through non-metric dimensional scaling (NMDS) and Curtis-Bray clustering analyses. Bold genera text denotes chemolithautotrophic sulfur oxidizing/disproportionating organisms.

| **Year** | **Family** | **Genus** | **Relative Abundance (%)** | **Relative Abundance Range** |
|---|---|---|---|---|
| | ***Thiovulaceae*** | ***Sulfuricurvum* sp.** | 37 ± 28 | 2 - 69 |
| | ***Thiobacillaceae*** | ***Thiobacillus* sp.** | 17 ± 20 | 1 - 56 |
| | ***Halothiobacillaceae*** | ***Halothiobacillus* sp.** | 12 ± 8 | 2 - 24 |
| | | ***Thiovirga* sp.** | 0.7 ± 0.9 | 0.01- 2 |
| **2015** | *Methylophilaceae* | *Methylotenera* sp. | 6 ± 4 | 4 - 14 |
| n = 8 | ***Chitinophagaceae*** | ***Sediminibacterium* sp.** | 5.9 ± 9 | 0.2 - 26 |
| | *Flavobacteriaceae* | *Lutibacter* sp. | 3.6 ± 2 | 2 - 6 |
| | ***Sulfurovaceae*** | ***Sulfurovum* sp.** | 3 ± 4 | 0.1 - 9 |
| | *Burkholderiaceae* | *Acidovorax* sp. | 1.7 ± 1 | 0.4 - 3 |
| | Other | | 13 | |
| | ***Halothiobacillaceae*** | ***Halothiobacillus* sp.** | 50 ± 21 | 12 - 78 |
| | | ***Thiovirga* sp.** | 0.5 ± 0.5 | 0.1 - 1.7 |
| | *Sphingomonadaceae* | *Sphingobium* sp. | 12 ± 13 | 3 - 42 |
| | | *Novosphingobium* sp. | 10 ± 6 | 3 - 22 |
| **2016/2017** | | *Sphingomonas* sp. | 8.9 ± 9 | 0.2 - 23 |
| **n = 13** | Chitinophagaceae | ***Sediminibacterium sp*.** | 1.5 ± 1.4 | 0.2 - 5.2 |
| | | *Hydrotalea* sp. | 1.0 ± 1.7 | 0.01 - 6 |
| | ***Thiovulaceae*** | ***Sulfuricurvum* sp.** | 1.8 ± 1.9 | <LOD - 5 |
| | ***Burkholderiaceae*** | ***Acidovorax* sp.** | 1 ± 0.8 | <LOD - 3 |
| | Other | | 13 | |
| | ***Thiovulaceae*** | ***Sulfuricurvum* sp.** | 17 ± 26 | <LOD - 68 |
| | *Chitinophagaceae* | ***Sediminibacterium sp*.** | 14 ± 16 | <LOD - 49 |
| | ***Halothiobacillaceae*** | ***Halothiobacillus* sp.** | 11 ± 11 | 0.1 - 34 |
| | | ***Thiovirga* sp.** | 5 ± 8 | <LOD - 24 |
| | *Burkholderiaceae* | ***Acidovorax* sp.** | 9 ± 11 | 0.3 - 32 |
| | | *Sphingomonas* sp. | 4 ± 6 | 0.1 - 19 |
| | *Sphingomonadaceae* | *Novosphingobium* sp. | 0.9 ± 1.3 | 0.05 - 1.3 |
| | | *Sphingobium* sp. | 0.5 ± 0.7 | <LOD - 2.4 |
| **2018** | *Sphingobacteriaceae* | *Pedobacter* sp. | 4 ± 8 | <LOD - 24 |
| **n = 9** | *Flavobacteriaceae* | *Flavobacterium* sp. | 2.0 ± 4 | <LOD - 10 |
| | | *Lutibacter* sp. | 0.8 ± 1 | 0.1 - 3.2 |
| | ***Thiobacillaceae*** | ***Thiobacillus* sp.** | 3.9 ± 6 | <LOD - 19 |
| | *Methylophilaceae* | *Methylotenera* sp. | 0.9 ± 2 | <LOD - 6.3 |
| | *Rhodocyclaceae* | *Dechloromonas* sp. | 1.1 ± 2 | <LOD - 6 |
| | *Mycoplasmataceae* | *Mycoplasma* sp. | 1.5 ± 5 | <LOD - 13 |
| | *Xanthobacteraceae* | *Xanthobacter* sp. | 1.5 ± 4 | <LOD - 11 |
| | *Caulobacteraceae* | *Caulobacter* sp. | 0.9 ± 2 | 0.1 - 4 |
| | Other | | 23 | |

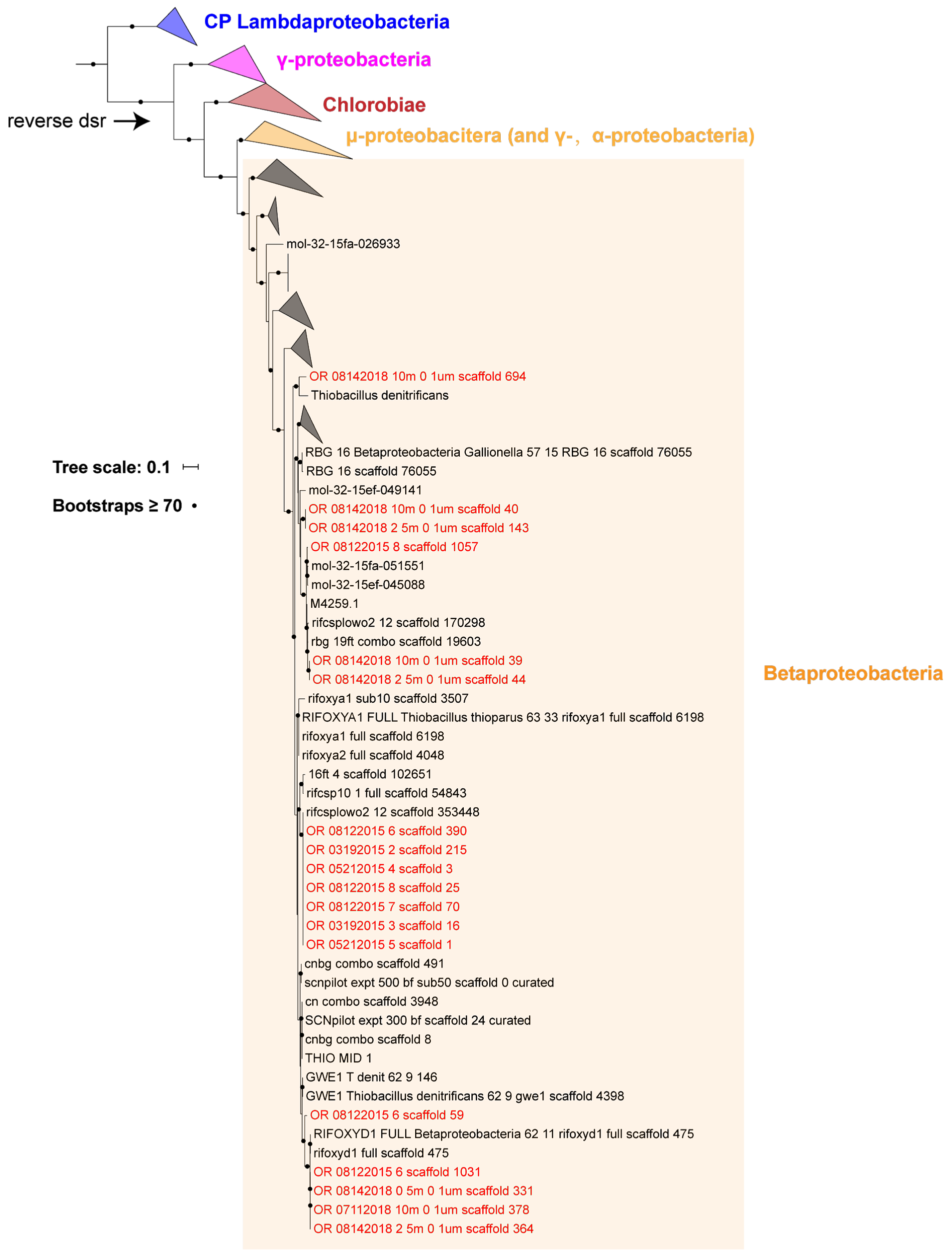


**Fig S1 | Concatenated DsrAB phylogenetic tree indicated the DsrAB identified in this study are likely responsible for sulfur oxidation (i.e., reverse dsr)**. The DsrAB sequences reported in this study are in red. See Methods for details of tree construction. Reference sequences are retrieved from [57].

### Table S5. Community level relative abundances of sulfur cycling nitrogen cycling, hydrogen cycling, carbon fixation, photosynthetic and aeronic respiration genes over time (2015-2018) in tailing impoundment waters,

| Gene | March 2015 0.5 m | March 2015 0.75 m | March 2015 21 m | May 2015 2.5 m | May 2015 3.5 m | August 2015 1 m | August 2015 2.8 m | August 2015 25.8 m | July 2016 1.5 m | July 2016 10 m | June 2017 2m | June 2017 10 m | July 2017 0.5 m | July 2017 2 m | July 2017 10 m | August 2017 0.5 m | August 2017 2m | August 2017 10 m | September 2017 10 m | November 2017 0.5 m | November 2017 10 m | May 2018 0.5 m | May 2018 2.8 m | May 2018 10 m | July 10 2018 0.5 m | July 10 2018 10 m | July 23 2018 0.5 m | August 2018 0.5 m | August 2018 2.5 m | Augut 2018 10 m |
|---|---|---|---|---|---|---|---|---|---|---|---|---|---|---|---|---|---|---|---|---|---|---|---|---|---|---|---|---|---|---|
| cbb3 | 0.01% | 0.01% | 0.01% | 0.01% | 0.00% | 0.00% | 0.00% | 0.00% | 0.00% | 0.00% | 0.00% | 0.00% | 0.00% | 0.00% | 0.00% | 0.00% | 0.00% | 0.00% | 0.00% | 0.00% | 0.00% | 0.00% | 0.00% | 0.00% | 0.00% | 0.00% | 0.00% | 0.00% | 0.00% | 0.00% |
| ccoN | 0.04% | 0.04% | 0.05% | 0.04% | 0.04% | 0.04% | 0.04% | 0.06% | 0.04% | 0.05% | 0.04% | 0.05% | 0.05% | 0.05% | 0.06% | 0.06% | 0.05% | 0.08% | 0.03% | 0.04% | 0.04% | 0.04% | 0.03% | 0.06% | 0.05% | 0.04% | 0.04% | 0.04% | 0.04% | 0.04% |
| ccoO | 0.03% | 0.04% | 0.03% | 0.02% | 0.04% | 0.04% | 0.04% | 0.04% | 0.03% | 0.03% | 0.04% | 0.04% | 0.04% | 0.04% | 0.04% | 0.04% | 0.03% | 0.04% | 0.03% | 0.04% | 0.04% | 0.03% | 0.03% | 0.02% | 0.03% | 0.04% | 0.04% | 0.04% | 0.04% | 0.04% |
| ccoP | 0.03% | 0.03% | 0.03% | 0.03% | 0.03% | 0.03% | 0.04% | 0.04% | 0.04% | 0.05% | 0.04% | 0.05% | 0.03% | 0.04% | 0.06% | 0.04% | 0.04% | 0.05% | 0.02% | 0.04% | 0.04% | 0.01% | 0.02% | 0.02% | 0.02% | 0.04% | 0.02% | 0.03% | 0.04% | 0.04% |
| CoxA | 0.02% | 0.02% | 0.02% | 0.02% | 0.02% | 0.02% | 0.02% | 0.01% | 0.01% | 0.01% | 0.02% | 0.01% | 0.01% | 0.02% | 0.01% | 0.01% | 0.01% | 0.01% | 0.01% | 0.02% | 0.02% | 0.02% | 0.02% | 0.00% | 0.02% | 0.01% | 0.02% | 0.02% | 0.01% | 0.01% |
| CoxB | 0.02% | 0.02% | 0.02% | 0.02% | 0.02% | 0.02% | 0.03% | 0.02% | 0.01% | 0.01% | 0.02% | 0.01% | 0.02% | 0.02% | 0.01% | 0.01% | 0.01% | 0.01% | 0.01% | 0.02% | 0.02% | 0.02% | 0.02% | 0.02% | 0.02% | 0.02% | 0.02% | 0.02% | 0.01% | 0.01% |
| cydA_Cyt_bd_oxida_I | 0.02% | 0.01% | 0.01% | 0.01% | 0.01% | 0.02% | 0.01% | 0.01% | 0.06% | 0.07% | 0.08% | 0.06% | 0.05% | 0.05% | 0.06% | 0.06% | 0.05% | 0.05% | 0.04% | 0.05% | 0.06% | 0.01% | 0.01% | 0.01% | 0.03% | 0.03% | 0.03% | 0.04% | 0.02% | 0.01% |
| cydB | 0.01% | 0.01% | 0.01% | 0.01% | 0.01% | 0.01% | 0.01% | 0.01% | 0.05% | 0.05% | 0.05% | 0.04% | 0.05% | 0.05% | 0.06% | 0.03% | 0.05% | 0.05% | 0.04% | 0.05% | 0.06% | 0.01% | 0.01% | 0.01% | 0.02% | 0.02% | 0.03% | 0.04% | 0.02% | 0.01% |
| CyoA | 0.00% | 0.00% | 0.00% | 0.00% | 0.00% | 0.00% | 0.00% | 0.00% | 0.01% | 0.01% | 0.01% | 0.01% | 0.01% | 0.01% | 0.00% | 0.00% | 0.00% | 0.00% | 0.01% | 0.01% | 0.01% | 0.00% | 0.00% | 0.00% | 0.00% | 0.00% | 0.00% | 0.00% | 0.00% | 0.00% |
| CyoB | 0.00% | 0.00% | 0.00% | 0.00% | 0.00% | 0.00% | 0.00% | 0.00% | 0.01% | 0.01% | 0.01% | 0.01% | 0.01% | 0.01% | 0.00% | 0.00% | 0.00% | 0.00% | 0.01% | 0.01% | 0.01% | 0.00% | 0.00% | 0.00% | 0.00% | 0.00% | 0.00% | 0.00% | 0.00% | 0.00% |
| CyoC | 0.00% | 0.00% | 0.00% | 0.00% | 0.00% | 0.00% | 0.00% | 0.00% | 0.01% | 0.01% | 0.01% | 0.01% | 0.01% | 0.01% | 0.00% | 0.00% | 0.00% | 0.00% | 0.01% | 0.01% | 0.01% | 0.00% | 0.00% | 0.00% | 0.00% | 0.00% | 0.00% | 0.00% | 0.00% | 0.00% |
| CyoD | 0.00% | 0.00% | 0.00% | 0.00% | 0.00% | 0.00% | 0.00% | 0.00% | 0.01% | 0.01% | 0.01% | 0.01% | 0.01% | 0.01% | 0.00% | 0.00% | 0.00% | 0.00% | 0.01% | 0.01% | 0.01% | 0.00% | 0.00% | 0.00% | 0.00% | 0.00% | 0.00% | 0.00% | 0.00% | 0.00% |
| pqq_xoxF_mxaF | 0.03% | 0.04% | 0.04% | 0.04% | 0.05% | 0.03% | 0.02% | 0.04% | 0.03% | 0.03% | 0.03% | 0.03% | 0.03% | 0.04% | 0.03% | 0.03% | 0.03% | 0.03% | 0.03% | 0.03% | 0.03% | 0.06% | 0.06% | 0.05% | 0.04% | 0.02% | 0.03% | 0.03% | 0.01% | 0.01% |
| aclA | 0.01% | 0.01% | 0.01% | 0.01% | 0.01% | 0.00% | 0.00% | 0.00% | 0.00% | 0.00% | 0.00% | 0.00% | 0.00% | 0.00% | 0.00% | 0.00% | 0.00% | 0.00% | 0.00% | 0.00% | 0.00% | 0.00% | 0.00% | 0.01% | 0.00% | 0.00% | 0.00% | 0.00% | 0.00% | 0.00% |
| aclB | 0.01% | 0.01% | 0.01% | 0.01% | 0.01% | 0.00% | 0.00% | 0.00% | 0.00% | 0.00% | 0.00% | 0.00% | 0.00% | 0.00% | 0.00% | 0.00% | 0.00% | 0.00% | 0.00% | 0.00% | 0.00% | 0.00% | 0.00% | 0.01% | 0.00% | 0.00% | 0.00% | 0.00% | 0.00% | 0.00% |
| rubisco_form_I | 0.01% | 0.01% | 0.01% | 0.00% | 0.01% | 0.01% | 0.01% | 0.01% | 0.02% | 0.02% | 0.01% | 0.02% | 0.01% | 0.01% | 0.02% | 0.02% | 0.02% | 0.02% | 0.01% | 0.01% | 0.00% | 0.00% | 0.00% | 0.00% | 0.01% | 0.01% | 0.01% | 0.02% | 0.02% | 0.02% |
| rubisco_form_II | 0.01% | 0.01% | 0.01% | 0.00% | 0.01% | 0.00% | 0.01% | 0.01% | 0.01% | 0.02% | 0.01% | 0.02% | 0.01% | 0.01% | 0.02% | 0.01% | 0.01% | 0.02% | 0.00% | 0.00% | 0.00% | 0.00% | 0.00% | 0.00% | 0.00% | 0.00% | 0.00% | 0.01% | 0.00% | 0.00% |
| nosZ | 0.01% | 0.01% | 0.01% | 0.00% | 0.00% | 0.01% | 0.00% | 0.00% | 0.00% | 0.00% | 0.00% | 0.00% | 0.00% | 0.00% | 0.00% | 0.00% | 0.00% | 0.00% | 0.00% | 0.00% | 0.00% | 0.00% | 0.00% | 0.00% | 0.01% | 0.01% | 0.01% | 0.01% | 0.01% | 0.01% |
| Hydrogenase_Group_1 | 0.01% | 0.01% | 0.01% | 0.02% | 0.01% | 0.00% | 0.00% | 0.01% | 0.00% | 0.00% | 0.00% | 0.00% | 0.00% | 0.00% | 0.00% | 0.00% | 0.00% | 0.00% | 0.01% | 0.00% | 0.00% | 0.00% | 0.00% | 0.01% | 0.00% | 0.00% | 0.00% | 0.00% | 0.01% | 0.01% |
| Hydrogenase_Group_2a | 0.00% | 0.00% | 0.00% | 0.00% | 0.00% | 0.00% | 0.00% | 0.00% | 0.00% | 0.00% | 0.00% | 0.00% | 0.00% | 0.00% | 0.00% | 0.00% | 0.00% | 0.00% | 0.00% | 0.00% | 0.00% | 0.00% | 0.00% | 0.00% | 0.00% | 0.00% | 0.00% | 0.00% | 0.00% | 0.00% |
| Hydrogenase_Group_2b | 0.01% | 0.00% | 0.00% | 0.00% | 0.00% | 0.00% | 0.00% | 0.00% | 0.00% | 0.00% | 0.00% | 0.00% | 0.00% | 0.00% | 0.00% | 0.00% | 0.00% | 0.00% | 0.01% | 0.00% | 0.00% | 0.00% | 0.00% | 0.00% | 0.00% | 0.00% | 0.00% | 0.00% | 0.00% | 0.00% |
| Hydrogenase_Group_3b | 0.00% | 0.00% | 0.00% | 0.00% | 0.00% | 0.00% | 0.00% | 0.01% | 0.01% | 0.01% | 0.01% | 0.02% | 0.00% | 0.01% | 0.02% | 0.01% | 0.01% | 0.02% | 0.00% | 0.00% | 0.00% | 0.00% | 0.00% | 0.00% | 0.00% | 0.00% | 0.00% | 0.00% | 0.00% | 0.01% |
| Hydrogenase_Group_3c | 0.00% | 0.00% | 0.00% | 0.00% | 0.00% | 0.00% | 0.00% | 0.00% | 0.00% | 0.00% | 0.00% | 0.00% | 0.00% | 0.00% | 0.00% | 0.00% | 0.00% | 0.00% | 0.00% | 0.00% | 0.00% | 0.00% | 0.00% | 0.00% | 0.00% | 0.00% | 0.00% | 0.00% | 0.00% | 0.00% |
| Hydrogenase_Group_3d | 0.01% | 0.00% | 0.01% | 0.00% | 0.00% | 0.00% | 0.00% | 0.00% | 0.00% | 0.00% | 0.00% | 0.00% | 0.00% | 0.00% | 0.00% | 0.00% | 0.00% | 0.00% | 0.00% | 0.00% | 0.00% | 0.00% | 0.00% | 0.00% | 0.00% | 0.00% | 0.00% | 0.00% | 0.00% | 0.00% |
| Hydrogenase_Group_4 | 0.00% | 0.00% | 0.00% | 0.00% | 0.00% | 0.00% | 0.00% | 0.00% | 0.00% | 0.00% | 0.00% | 0.00% | 0.00% | 0.00% | 0.00% | 0.00% | 0.00% | 0.00% | 0.00% | 0.00% | 0.00% | 0.00% | 0.00% | 0.00% | 0.00% | 0.00% | 0.00% | 0.00% | 0.00% | 0.00% |
| cysA | 0.01% | 0.01% | 0.01% | 0.01% | 0.01% | 0.00% | 0.00% | 0.01% | 0.00% | 0.00% | 0.00% | 0.00% | 0.00% | 0.00% | 0.00% | 0.00% | 0.00% | 0.00% | 0.00% | 0.00% | 0.00% | 0.02% | 0.01% | 0.00% | 0.01% | 0.00% | 0.00% | 0.00% | 0.00% | 0.01% |
| cysD | 0.01% | 0.02% | 0.01% | 0.01% | 0.01% | 0.01% | 0.01% | 0.00% | 0.01% | 0.01% | 0.01% | 0.01% | 0.01% | 0.01% | 0.01% | 0.01% | 0.01% | 0.01% | 0.01% | 0.02% | 0.02% | 0.04% | 0.04% | 0.00% | 0.01% | 0.01% | 0.02% | 0.01% | 0.01% | 0.01% |
| cysI | 0.01% | 0.01% | 0.01% | 0.01% | 0.01% | 0.01% | 0.01% | 0.01% | 0.02% | 0.01% | 0.01% | 0.01% | 0.01% | 0.01% | 0.01% | 0.01% | 0.01% | 0.00% | 0.01% | 0.02% | 0.02% | 0.02% | 0.02% | 0.00% | 0.01% | 0.01% | 0.01% | 0.01% | 0.00% | 0.00% |
| cysJ | 0.00% | 0.00% | 0.00% | 0.00% | 0.00% | 0.00% | 0.00% | 0.00% | 0.00% | 0.00% | 0.00% | 0.00% | 0.00% | 0.00% | 0.00% | 0.00% | 0.00% | 0.00% | 0.00% | 0.00% | 0.00% | 0.00% | 0.00% | 0.00% | 0.00% | 0.00% | 0.00% | 0.00% | 0.00% | 0.00% |
| cysN_cysC | 0.00% | 0.00% | 0.00% | 0.00% | 0.00% | 0.00% | 0.00% | 0.00% | 0.01% | 0.00% | 0.01% | 0.01% | 0.01% | 0.01% | 0.00% | 0.00% | 0.00% | 0.00% | 0.01% | 0.01% | 0.01% | 0.02% | 0.02% | 0.00% | 0.01% | 0.00% | 0.01% | 0.00% | 0.00% | 0.00% |
| cysP | 0.00% | 0.00% | 0.00% | 0.00% | 0.00% | 0.00% | 0.00% | 0.00% | 0.00% | 0.00% | 0.00% | 0.00% | 0.00% | 0.00% | 0.00% | 0.00% | 0.00% | 0.00% | 0.00% | 0.00% | 0.00% | 0.00% | 0.00% | 0.00% | 0.00% | 0.00% | 0.00% | 0.00% | 0.00% | 0.00% |
| cysU | 0.01% | 0.00% | 0.01% | 0.01% | 0.01% | 0.00% | 0.00% | 0.00% | 0.00% | 0.00% | 0.00% | 0.00% | 0.00% | 0.00% | 0.00% | 0.00% | 0.00% | 0.00% | 0.00% | 0.00% | 0.00% | 0.01% | 0.00% | 0.00% | 0.00% | 0.00% | 0.00% | 0.00% | 0.00% | 0.00% |
| cysW | 0.01% | 0.00% | 0.00% | 0.01% | 0.01% | 0.00% | 0.00% | 0.00% | 0.00% | 0.00% | 0.00% | 0.00% | 0.00% | 0.00% | 0.00% | 0.00% | 0.00% | 0.00% | 0.00% | 0.00% | 0.00% | 0.02% | 0.00% | 0.00% | 0.00% | 0.00% | 0.00% | 0.00% | 0.00% | 0.00% |
| doxD | 0.00% | 0.00% | 0.00% | 0.00% | 0.00% | 0.00% | 0.00% | 0.00% | 0.00% | 0.00% | 0.00% | 0.00% | 0.00% | 0.00% | 0.00% | 0.00% | 0.00% | 0.00% | 0.00% | 0.00% | 0.00% | 0.00% | 0.00% | 0.00% | 0.00% | 0.00% | 0.00% | 0.00% | 0.00% | 0.00% |
| hdrA2 | 0.00% | 0.00% | 0.00% | 0.00% | 0.00% | 0.00% | 0.00% | 0.00% | 0.00% | 0.00% | 0.00% | 0.00% | 0.00% | 0.00% | 0.00% | 0.00% | 0.00% | 0.00% | 0.00% | 0.00% | 0.00% | 0.00% | 0.00% | 0.00% | 0.00% | 0.00% | 0.00% | 0.00% | 0.00% | 0.00% |
| hdrB2 | 0.00% | 0.00% | 0.00% | 0.00% | 0.00% | 0.00% | 0.00% | 0.00% | 0.00% | 0.00% | 0.00% | 0.00% | 0.00% | 0.00% | 0.00% | 0.00% | 0.00% | 0.00% | 0.00% | 0.00% | 0.00% | 0.00% | 0.00% | 0.00% | 0.00% | 0.00% | 0.00% | 0.00% | 0.00% | 0.00% |
| hdrC2 | 0.00% | 0.00% | 0.00% | 0.00% | 0.00% | 0.00% | 0.00% | 0.00% | 0.00% | 0.00% | 0.00% | 0.00% | 0.00% | 0.00% | 0.00% | 0.00% | 0.00% | 0.00% | 0.00% | 0.00% | 0.00% | 0.00% | 0.00% | 0.00% | 0.00% | 0.00% | 0.00% | 0.00% | 0.00% | 0.00% |
| psrA | 0.00% | 0.00% | 0.00% | 0.00% | 0.00% | 0.00% | 0.00% | 0.00% | 0.00% | 0.00% | 0.00% | 0.00% | 0.00% | 0.00% | 0.00% | 0.00% | 0.00% | 0.00% | 0.00% | 0.00% | 0.00% | 0.00% | 0.00% | 0.00% | 0.00% | 0.00% | 0.00% | 0.00% | 0.00% | 0.00% |
| qmoA | 0.00% | 0.00% | 0.00% | 0.00% | 0.00% | 0.00% | 0.01% | 0.01% | 0.00% | 0.00% | 0.00% | 0.00% | 0.00% | 0.00% | 0.00% | 0.00% | 0.00% | 0.00% | 0.00% | 0.00% | 0.00% | 0.00% | 0.00% | 0.00% | 0.00% | 0.00% | 0.00% | 0.00% | 0.01% | 0.01% |
| qmoB | 0.00% | 0.00% | 0.00% | 0.00% | 0.00% | 0.00% | 0.01% | 0.01% | 0.00% | 0.00% | 0.00% | 0.00% | 0.00% | 0.00% | 0.00% | 0.00% | 0.00% | 0.00% | 0.00% | 0.00% | 0.00% | 0.00% | 0.00% | 0.00% | 0.00% | 0.00% | 0.00% | 0.00% | 0.01% | 0.01% |
| qmoC | 0.00% | 0.00% | 0.00% | 0.00% | 0.00% | 0.00% | 0.00% | 0.00% | 0.00% | 0.00% | 0.00% | 0.00% | 0.00% | 0.00% | 0.00% | 0.00% | 0.00% | 0.00% | 0.00% | 0.00% | 0.00% | 0.00% | 0.00% | 0.00% | 0.00% | 0.00% | 0.00% | 0.00% | 0.00% | 0.00% |
| sir | 0.00% | 0.00% | 0.00% | 0.00% | 0.00% | 0.00% | 0.00% | 0.00% | 0.00% | 0.00% | 0.00% | 0.00% | 0.00% | 0.00% | 0.00% | 0.00% | 0.00% | 0.00% | 0.00% | 0.00% | 0.00% | 0.00% | 0.00% | 0.00% | 0.00% | 0.00% | 0.00% | 0.00% | 0.00% | 0.00% |
| soeA | 0.00% | 0.00% | 0.00% | 0.00% | 0.00% | 0.01% | 0.01% | 0.01% | 0.00% | 0.00% | 0.00% | 0.00% | 0.00% | 0.00% | 0.00% | 0.00% | 0.00% | 0.00% | 0.00% | 0.00% | 0.00% | 0.00% | 0.00% | 0.00% | 0.00% | 0.00% | 0.00% | 0.00% | 0.01% | 0.01% |
| soeB | 0.00% | 0.00% | 0.00% | 0.00% | 0.00% | 0.01% | 0.01% | 0.01% | 0.00% | 0.00% | 0.00% | 0.00% | 0.00% | 0.00% | 0.00% | 0.00% | 0.00% | 0.00% | 0.00% | 0.00% | 0.00% | 0.00% | 0.00% | 0.00% | 0.00% | 0.00% | 0.00% | 0.00% | 0.01% | 0.00% |
| soeC | 0.00% | 0.00% | 0.00% | 0.00% | 0.00% | 0.01% | 0.01% | 0.01% | 0.00% | 0.00% | 0.00% | 0.00% | 0.00% | 0.00% | 0.00% | 0.00% | 0.00% | 0.00% | 0.00% | 0.00% | 0.00% | 0.00% | 0.00% | 0.00% | 0.00% | 0.00% | 0.00% | 0.00% | 0.01% | 0.01% |
| SoxD | 0.02% | 0.02% | 0.02% | 0.01% | 0.01% | 0.00% | 0.00% | 0.00% | 0.02% | 0.02% | 0.00% | 0.00% | 0.01% | 0.01% | 0.02% | 0.00% | 0.00% | 0.00% | 0.01% | 0.01% | 0.00% | 0.00% | 0.00% | 0.00% | 0.00% | 0.00% | 0.00% | 0.00% | 0.00% | 0.00% |
| sqr | 0.03% | 0.04% | 0.04% | 0.05% | 0.05% | 0.02% | 0.04% | 0.06% | 0.04% | 0.04% | 0.04% | 0.06% | 0.03% | 0.04% | 0.06% | 0.04% | 0.02% | 0.04% | 0.02% | 0.04% | 0.03% | 0.04% | 0.04% | 0.00% | 0.04% | 0.04% | 0.03% | 0.02% | 0.03% | 0.03% |
| tsdA | 0.01% | 0.01% | 0.01% | 0.01% | 0.01% | 0.01% | 0.01% | 0.01% | 0.02% | 0.02% | 0.02% | 0.02% | 0.02% | 0.01% | 0.02% | 0.02% | 0.02% | 0.02% | 0.02% | 0.02% | 0.02% | 0.00% | 0.00% | 0.00% | 0.02% | 0.03% | 0.02% | 0.02% | 0.02% | 0.01% |
| TST | 0.02% | 0.03% | 0.03% | 0.02% | 0.02% | 0.01% | 0.02% | 0.03% | 0.03% | 0.03% | 0.02% | 0.03% | 0.01% | 0.02% | 0.03% | 0.02% | 0.02% | 0.02% | 0.02% | 0.02% | 0.02% | 0.02% | 0.02% | 0.01% | 0.02% | 0.01% | 0.01% | 0.01% | 0.02% | 0.02% |
| ttrA | 0.00% | 0.01% | 0.01% | 0.01% | 0.01% | 0.00% | 0.00% | 0.02% | 0.00% | 0.00% | 0.00% | 0.00% | 0.00% | 0.00% | 0.00% | 0.00% | 0.00% | 0.00% | 0.00% | 0.00% | 0.00% | 0.02% | 0.01% | 0.00% | 0.01% | 0.00% | 0.00% | 0.00% | 0.00% | 0.01% |
| ttrB | 0.00% | 0.01% | 0.01% | 0.01% | 0.01% | 0.00% | 0.00% | 0.01% | 0.00% | 0.00% | 0.00% | 0.00% | 0.00% | 0.00% | 0.00% | 0.00% | 0.00% | 0.00% | 0.00% | 0.00% | 0.00% | 0.02% | 0.01% | 0.00% | 0.01% | 0.00% | 0.00% | 0.00% | 0.00% | 0.01% |
| ttrC | 0.00% | 0.01% | 0.01% | 0.01% | 0.01% | 0.00% | 0.00% | 0.01% | 0.00% | 0.00% | 0.00% | 0.00% | 0.00% | 0.00% | 0.00% | 0.00% | 0.00% | 0.00% | 0.00% | 0.00% | 0.00% | 0.02% | 0.01% | 0.00% | 0.01% | 0.00% | 0.00% | 0.00% | 0.00% | 0.01% |
| 1.3.1.33_protochlorophyllide_reductase | 0.00% | 0.00% | 0.00% | 0.00% | 0.00% | 0.00% | 0.00% | 0.00% | 0.00% | 0.00% | 0.00% | 0.00% | 0.00% | 0.00% | 0.00% | 0.00% | 0.00% | 0.00% | 0.01% | 0.00% | 0.00% | 0.00% | 0.00% | 0.00% | 0.01% | 0.01% | 0.01% | 0.01% | 0.00% | 0.00% |
| bchC | 0.00% | 0.00% | 0.00% | 0.00% | 0.00% | 0.00% | 0.00% | 0.00% | 0.00% | 0.00% | 0.01% | 0.01% | 0.01% | 0.01% | 0.00% | 0.00% | 0.00% | 0.00% | 0.01% | 0.01% | 0.01% | 0.00% | 0.00% | 0.00% | 0.00% | 0.00% | 0.01% | 0.00% | 0.00% | 0.00% |
| bchF | 0.00% | 0.00% | 0.00% | 0.00% | 0.00% | 0.00% | 0.00% | 0.00% | 0.00% | 0.00% | 0.01% | 0.01% | 0.01% | 0.01% | 0.00% | 0.00% | 0.00% | 0.00% | 0.01% | 0.01% | 0.01% | 0.00% | 0.00% | 0.00% | 0.00% | 0.00% | 0.01% | 0.00% | 0.00% | 0.00% |
| bchJ_divinyl_protochlorophyllide | 0.00% | 0.00% | 0.00% | 0.00% | 0.00% | 0.00% | 0.00% | 0.00% | 0.00% | 0.00% | 0.00% | 0.00% | 0.00% | 0.00% | 0.00% | 0.00% | 0.00% | 0.00% | 0.01% | 0.00% | 0.00% | 0.00% | 0.00% | 0.00% | 0.00% | 0.00% | 0.00% | 0.00% | 0.00% | 0.00% |
| bchX | 0.00% | 0.00% | 0.00% | 0.00% | 0.00% | 0.00% | 0.00% | 0.00% | 0.00% | 0.00% | 0.01% | 0.01% | 0.01% | 0.00% | 0.00% | 0.00% | 0.00% | 0.00% | 0.01% | 0.01% | 0.01% | 0.00% | 0.00% | 0.00% | 0.00% | 0.00% | 0.01% | 0.00% | 0.00% | 0.00% |
| bchY | 0.00% | 0.00% | 0.00% | 0.00% | 0.00% | 0.00% | 0.00% | 0.00% | 0.00% | 0.00% | 0.01% | 0.01% | 0.01% | 0.01% | 0.00% | 0.00% | 0.00% | 0.00% | 0.01% | 0.01% | 0.01% | 0.00% | 0.00% | 0.00% | 0.00% | 0.00% | 0.01% | 0.00% | 0.00% | 0.00% |
| bchZ | 0.00% | 0.00% | 0.00% | 0.00% | 0.00% | 0.00% | 0.00% | 0.00% | 0.00% | 0.00% | 0.01% | 0.01% | 0.01% | 0.01% | 0.00% | 0.00% | 0.00% | 0.00% | 0.01% | 0.01% | 0.01% | 0.00% | 0.00% | 0.00% | 0.00% | 0.00% | 0.01% | 0.00% | 0.00% | 0.00% |
| chlB_light-independent_protochlorophyllide_reductase_subunit_B | 0.00% | 0.00% | 0.00% | 0.00% | 0.00% | 0.01% | 0.00% | 0.00% | 0.00% | 0.00% | 0.01% | 0.01% | 0.01% | 0.01% | 0.00% | 0.00% | 0.00% | 0.00% | 0.01% | 0.01% | 0.01% | 0.00% | 0.00% | 0.00% | 0.00% | 0.00% | 0.01% | 0.00% | 0.00% | 0.00% |
| chlG_bchG_chlorophyll_bacteriochlorophyll_a_synthase | 0.00% | 0.00% | 0.00% | 0.00% | 0.00% | 0.00% | 0.00% | 0.00% | 0.00% | 0.00% | 0.01% | 0.01% | 0.01% | 0.01% | 0.00% | 0.00% | 0.00% | 0.00% | 0.01% | 0.01% | 0.01% | 0.00% | 0.00% | 0.00% | 0.00% | 0.00% | 0.01% | 0.00% | 0.00% | 0.00% |
| chlL_light-independent_protochlorophyllide_reductase_subunit_L | 0.00% | 0.00% | 0.00% | 0.00% | 0.00% | 0.00% | 0.00% | 0.00% | 0.00% | 0.00% | 0.01% | 0.01% | 0.01% | 0.01% | 0.00% | 0.00% | 0.00% | 0.00% | 0.01% | 0.01% | 0.01% | 0.00% | 0.00% | 0.00% | 0.00% | 0.00% | 0.01% | 0.00% | 0.00% | 0.00% |
| chlN_light-independent_protochlorophyllide_reductase_subunit_N | 0.00% | 0.00% | 0.00% | 0.00% | 0.00% | 0.01% | 0.00% | 0.00% | 0.00% | 0.00% | 0.01% | 0.01% | 0.01% | 0.01% | 0.00% | 0.00% | 0.00% | 0.00% | 0.01% | 0.01% | 0.01% | 0.00% | 0.00% | 0.00% | 0.00% | 0.00% | 0.01% | 0.00% | 0.00% | 0.00% |
| chlP_bchP_geranylgeranyl_diphosphate_geranylgeranyl-bacteriochlorophyllide | 0.00% | 0.00% | 0.00% | 0.00% | 0.00% | 0.00% | 0.00% | 0.00% | 0.00% | 0.00% | 0.01% | 0.01% | 0.01% | 0.00% | 0.00% | 0.00% | 0.00% | 0.00% | 0.01% | 0.01% | 0.01% | 0.00% | 0.00% | 0.00% | 0.00% | 0.00% | 0.01% | 0.00% | 0.00% | 0.00% |
| pucA_light-harvesting_protein_B-800-850_alpha_chain | 0.00% | 0.00% | 0.00% | 0.00% | 0.00% | 0.00% | 0.00% | 0.00% | 0.00% | 0.00% | 0.00% | 0.00% | 0.00% | 0.00% | 0.00% | 0.00% | 0.00% | 0.00% | 0.00% | 0.00% | 0.00% | 0.00% | 0.00% | 0.00% | 0.00% | 0.00% | 0.00% | 0.00% | 0.00% | 0.00% |
| pucB_light-harvesting_protein_B-800-850_beta_chain | 0.00% | 0.00% | 0.00% | 0.00% | 0.00% | 0.00% | 0.00% | 0.00% | 0.00% | 0.00% | 0.00% | 0.00% | 0.00% | 0.00% | 0.00% | 0.00% | 0.00% | 0.00% | 0.00% | 0.00% | 0.00% | 0.00% | 0.00% | 0.00% | 0.00% | 0.00% | 0.00% | 0.00% | 0.00% | 0.00% |
| pufA_light-harvesting_complex_1_alpha_chain | 0.00% | 0.00% | 0.00% | 0.00% | 0.00% | 0.00% | 0.00% | 0.00% | 0.00% | 0.00% | 0.01% | 0.01% | 0.01% | 0.01% | 0.00% | 0.00% | 0.00% | 0.00% | 0.01% | 0.01% | 0.01% | 0.00% | 0.00% | 0.00% | 0.00% | 0.00% | 0.01% | 0.00% | 0.00% | 0.00% |
| pufB_light-harvesting_complex_1_beta_chain | 0.00% | 0.00% | 0.00% | 0.00% | 0.00% | 0.00% | 0.00% | 0.00% | 0.00% | 0.00% | 0.01% | 0.01% | 0.01% | 0.01% | 0.00% | 0.00% | 0.00% | 0.00% | 0.01% | 0.01% | 0.01% | 0.00% | 0.00% | 0.00% | 0.00% | 0.00% | 0.01% | 0.00% | 0.00% | 0.00% |
| pufL_photosynthetic_reaction_center_L_subunit | 0.00% | 0.00% | 0.00% | 0.00% | 0.00% | 0.00% | 0.00% | 0.00% | 0.00% | 0.00% | 0.01% | 0.01% | 0.01% | 0.01% | 0.00% | 0.00% | 0.00% | 0.00% | 0.01% | 0.01% | 0.01% | 0.00% | 0.00% | 0.00% | 0.00% | 0.00% | 0.01% | 0.00% | 0.00% | 0.00% |
| pufM_photosynthetic_reaction_center_M_subunit | 0.00% | 0.00% | 0.00% | 0.00% | 0.00% | 0.00% | 0.00% | 0.00% | 0.00% | 0.00% | 0.01% | 0.01% | 0.01% | 0.01% | 0.00% | 0.00% | 0.00% | 0.00% | 0.01% | 0.01% | 0.01% | 0.00% | 0.00% | 0.00% | 0.00% | 0.00% | 0.01% | 0.00% | 0.00% | 0.00% |
| pufX_photosynthetic_reaction_center_PufX_protein | 0.00% | 0.00% | 0.00% | 0.00% | 0.00% | 0.00% | 0.00% | 0.00% | 0.00% | 0.00% | 0.00% | 0.00% | 0.00% | 0.00% | 0.00% | 0.00% | 0.00% | 0.00% | 0.00% | 0.00% | 0.00% | 0.00% | 0.00% | 0.00% | 0.00% | 0.00% | 0.00% | 0.00% | 0.00% | 0.00% |
| puhA_photosynthetic_reaction_center_H_subunit | 0.00% | 0.00% | 0.00% | 0.00% | 0.00% | 0.01% | 0.00% | 0.00% | 0.00% | 0.00% | 0.01% | 0.01% | 0.01% | 0.01% | 0.00% | 0.00% | 0.00% | 0.00% | 0.02% | 0.01% | 0.01% | 0.00% | 0.00% | 0.00% | 0.00% | 0.01% | 0.01% | 0.00% | 0.00% | 0.01% |
| bacterial_rhodopsin | 0.00% | 0.00% | 0.00% | 0.00% | 0.00% | 0.01% | 0.01% | 0.00% | 0.00% | 0.00% | 0.00% | 0.00% | 0.01% | 0.00% | 0.00% | 0.00% | 0.00% | 0.00% | 0.01% | 0.00% | 0.00% | 0.00% | 0.00% | 0.00% | 0.01% | 0.01% | 0.01% | 0.01% | 0.01% | 0.00% |
| cynS | 0.02% | 0.02% | 0.02% | 0.02% | 0.03% | 0.01% | 0.01% | 0.02% | 0.01% | 0.01% | 0.00% | 0.00% | 0.00% | 0.01% | 0.01% | 0.01% | 0.01% | 0.01% | 0.00% | 0.00% | 0.00% | 0.02% | 0.02% | 0.03% | 0.01% | 0.01% | 0.01% | 0.00% | 0.01% | 0.01% |
| napA | 0.01% | 0.01% | 0.01% | 0.02% | 0.01% | 0.00% | 0.00% | 0.00% | 0.00% | 0.00% | 0.00% | 0.00% | 0.00% | 0.00% | 0.00% | 0.00% | 0.00% | 0.00% | 0.00% | 0.00% | 0.00% | 0.00% | 0.00% | 0.02% | 0.00% | 0.00% | 0.00% | 0.00% | 0.00% | 0.00% |
| napB | 0.02% | 0.01% | 0.01% | 0.02% | 0.01% | 0.00% | 0.00% | 0.00% | 0.00% | 0.00% | 0.00% | 0.00% | 0.00% | 0.00% | 0.00% | 0.00% | 0.00% | 0.00% | 0.00% | 0.00% | 0.00% | 0.00% | 0.00% | 0.02% | 0.00% | 0.00% | 0.00% | 0.00% | 0.00% | 0.00% |
| narG | 0.01% | 0.01% | 0.02% | 0.01% | 0.01% | 0.01% | 0.01% | 0.02% | 0.01% | 0.01% | 0.01% | 0.01% | 0.00% | 0.02% | 0.01% | 0.01% | 0.01% | 0.01% | 0.00% | 0.02% | 0.04% | 0.04% | 0.03% | 0.03% | 0.02% | 0.01% | 0.01% | 0.00% | 0.01% | 0.01% |
| narH | 0.01% | 0.01% | 0.00% | 0.01% | 0.01% | 0.01% | 0.01% | 0.01% | 0.01% | 0.01% | 0.01% | 0.01% | 0.00% | 0.02% | 0.01% | 0.01% | 0.01% | 0.01% | 0.00% | 0.02% | 0.04% | 0.02% | 0.02% | 0.01% | 0.01% | 0.01% | 0.01% | 0.00% | 0.01% | 0.01% |
| narI | 0.01% | 0.01% | 0.01% | 0.01% | 0.01% | 0.01% | 0.01% | 0.01% | 0.01% | 0.01% | 0.01% | 0.01% | 0.00% | 0.02% | 0.01% | 0.01% | 0.01% | 0.01% | 0.00% | 0.02% | 0.04% | 0.02% | 0.02% | 0.02% | 0.01% | 0.01% | 0.01% | 0.00% | 0.01% | 0.01% |
| narJ | 0.01% | 0.01% | 0.01% | 0.01% | 0.01% | 0.01% | 0.01% | 0.02% | 0.01% | 0.01% | 0.01% | 0.01% | 0.00% | 0.01% | 0.01% | 0.01% | 0.01% | 0.01% | 0.00% | 0.02% | 0.04% | 0.00% | 0.00% | 0.00% | 0.00% | 0.00% | 0.00% | 0.00% | 0.00% | 0.00% |
| nirB | 0.02% | 0.02% | 0.02% | 0.01% | 0.02% | 0.01% | 0.02% | 0.02% | 0.02% | 0.03% | 0.02% | 0.02% | 0.01% | 0.02% | 0.03% | 0.02% | 0.03% | 0.03% | 0.01% | 0.01% | 0.01% | 0.02% | 0.02% | 0.00% | 0.00% | 0.01% | 0.01% | 0.01% | 0.01% | 0.01% |
| nirK | 0.00% | 0.00% | 0.00% | 0.00% | 0.00% | 0.00% | 0.00% | 0.00% | 0.00% | 0.00% | 0.00% | 0.00% | 0.00% | 0.00% | 0.00% | 0.00% | 0.00% | 0.00% | 0.01% | 0.00% | 0.00% | 0.00% | 0.00% | 0.00% | 0.01% | 0.01% | 0.01% | 0.01% | 0.01% | 0.01% |
| nirS | 0.01% | 0.01% | 0.01% | 0.01% | 0.01% | 0.01% | 0.00% | 0.01% | 0.00% | 0.00% | 0.00% | 0.00% | 0.00% | 0.00% | 0.00% | 0.00% | 0.00% | 0.00% | 0.00% | 0.00% | 0.00% | 0.02% | 0.02% | 0.02% | 0.01% | 0.01% | 0.00% | 0.00% | 0.01% | 0.01% |
| norB | 0.02% | 0.02% | 0.02% | 0.02% | 0.02% | 0.01% | 0.02% | 0.03% | 0.03% | 0.03% | 0.03% | 0.03% | 0.02% | 0.03% | 0.03% | 0.03% | 0.03% | 0.03% | 0.01% | 0.02% | 0.02% | 0.02% | 0.02% | 0.02% | 0.02% | 0.02% | 0.02% | 0.02% | 0.02% | 0.02% |
| norC | 0.02% | 0.02% | 0.02% | 0.01% | 0.02% | 0.01% | 0.01% | 0.03% | 0.02% | 0.02% | 0.01% | 0.02% | 0.01% | 0.02% | 0.03% | 0.02% | 0.02% | 0.03% | 0.00% | 0.00% | 0.00% | 0.02% | 0.02% | 0.02% | 0.02% | 0.02% | 0.02% | 0.01% | 0.02% | 0.02% |
| nrfH | 0.00% | 0.00% | 0.00% | 0.00% | 0.00% | 0.00% | 0.00% | 0.00% | 0.00% | 0.00% | 0.00% | 0.00% | 0.00% | 0.00% | 0.00% | 0.00% | 0.00% | 0.00% | 0.00% | 0.00% | 0.00% | 0.00% | 0.00% | 0.00% | 0.00% | 0.00% | 0.00% | 0.00% | 0.00% | 0.00% |
| nxrA | 0.00% | 0.00% | 0.00% | 0.00% | 0.00% | 0.00% | 0.00% | 0.00% | 0.00% | 0.00% | 0.00% | 0.00% | 0.00% | 0.00% | 0.00% | 0.00% | 0.00% | 0.00% | 0.00% | 0.00% | 0.00% | 0.00% | 0.00% | 0.00% | 0.00% | 0.00% | 0.00% | 0.00% | 0.00% | 0.00% |
| nxrB | 0.00% | 0.00% | 0.00% | 0.00% | 0.00% | 0.00% | 0.00% | 0.00% | 0.00% | 0.00% | 0.00% | 0.00% | 0.00% | 0.00% | 0.00% | 0.00% | 0.00% | 0.00% | 0.00% | 0.00% | 0.00% | 0.00% | 0.00% | 0.00% | 0.00% | 0.00% | 0.00% | 0.00% | 0.00% | 0.00% |
| nifB | 0.00% | 0.00% | 0.00% | 0.01% | 0.01% | 0.00% | 0.00% | 0.00% | 0.00% | 0.00% | 0.00% | 0.00% | 0.00% | 0.00% | 0.00% | 0.00% | 0.00% | 0.00% | 0.00% | 0.00% | 0.00% | 0.00% | 0.00% | 0.00% | 0.00% | 0.00% | 0.00% | 0.00% | 0.00% | 0.00% |
| nifD | 0.00% | 0.00% | 0.00% | 0.01% | 0.01% | 0.00% | 0.00% | 0.00% | 0.00% | 0.00% | 0.00% | 0.00% | 0.00% | 0.00% | 0.00% | 0.00% | 0.00% | 0.00% | 0.00% | 0.00% | 0.00% | 0.00% | 0.00% | 0.00% | 0.00% | 0.00% | 0.00% | 0.00% | 0.00% | 0.00% |
| nifE | 0.00% | 0.00% | 0.00% | 0.01% | 0.01% | 0.00% | 0.00% | 0.01% | 0.00% | 0.00% | 0.00% | 0.00% | 0.00% | 0.00% | 0.00% | 0.00% | 0.00% | 0.00% | 0.00% | 0.00% | 0.00% | 0.00% | 0.00% | 0.01% | 0.00% | 0.00% | 0.00% | 0.01% | 0.00% | 0.00% |
| nifH | 0.00% | 0.00% | 0.00% | 0.01% | 0.01% | 0.00% | 0.00% | 0.00% | 0.00% | 0.00% | 0.00% | 0.00% | 0.00% | 0.00% | 0.00% | 0.00% | 0.00% | 0.00% | 0.00% | 0.00% | 0.00% | 0.00% | 0.00% | 0.00% | 0.00% | 0.00% | 0.00% | 0.00% | 0.00% | 0.00% |
| nifK | 0.00% | 0.00% | 0.00% | 0.01% | 0.01% | 0.00% | 0.00% | 0.00% | 0.00% | 0.00% | 0.00% | 0.00% | 0.00% | 0.00% | 0.00% | 0.00% | 0.00% | 0.00% | 0.00% | 0.00% | 0.00% | 0.00% | 0.00% | 0.00% | 0.00% | 0.00% | 0.00% | 0.00% | 0.00% | 0.00% |
| nifQ | 0.00% | 0.00% | 0.00% | 0.01% | 0.01% | 0.00% | 0.00% | 0.00% | 0.00% | 0.00% | 0.00% | 0.00% | 0.00% | 0.00% | 0.00% | 0.00% | 0.00% | 0.00% | 0.00% | 0.00% | 0.00% | 0.00% | 0.00% | 0.01% | 0.00% | 0.00% | 0.00% | 0.00% | 0.00% | 0.00% |
| nifT | 0.00% | 0.00% | 0.00% | 0.01% | 0.01% | 0.00% | 0.00% | 0.00% | 0.00% | 0.00% | 0.00% | 0.00% | 0.00% | 0.00% | 0.00% | 0.00% | 0.00% | 0.00% | 0.00% | 0.00% | 0.00% | 0.00% | 0.00% | 0.01% | 0.00% | 0.00% | 0.00% | 0.00% | 0.00% | 0.00% |
| nifX | 0.01% | 0.01% | 0.01% | 0.02% | 0.02% | 0.00% | 0.00% | 0.01% | 0.00% | 0.00% | 0.00% | 0.00% | 0.00% | 0.00% | 0.00% | 0.00% | 0.00% | 0.00% | 0.00% | 0.00% | 0.00% | 0.00% | 0.00% | 0.01% | 0.00% | 0.00% | 0.00% | 0.00% | 0.00% | 0.00% |
| nifZ | 0.02% | 0.01% | 0.01% | 0.04% | 0.03% | 0.01% | 0.01% | 0.02% | 0.01% | 0.01% | 0.01% | 0.01% | 0.00% | 0.01% | 0.01% | 0.01% | 0.01% | 0.01% | 0.00% | 0.03% | 0.04% | 0.00% | 0.00% | 0.02% | 0.00% | 0.01% | 0.00% | 0.01% | 0.02% | 0.02% |
| nosD | 0.02% | 0.01% | 0.02% | 0.01% | 0.01% | 0.01% | 0.01% | 0.02% | 0.00% | 0.00% | 0.00% | 0.00% | 0.00% | 0.00% | 0.01% | 0.01% | 0.01% | 0.01% | 0.00% | 0.00% | 0.00% | 0.02% | 0.02% | 0.02% | 0.01% | 0.01% | 0.01% | 0.01% | 0.01% | 0.02% |
| aprA | 0.00% | 0.00% | 0.00% | 0.00% | 0.01% | 0.01% | 0.02% | 0.02% | 0.00% | 0.00% | 0.00% | 0.00% | 0.00% | 0.00% | 0.00% | 0.00% | 0.00% | 0.00% | 0.00% | 0.00% | 0.00% | 0.00% | 0.00% | 0.00% | 0.00% | 0.00% | 0.00% | 0.00% | 0.01% | 0.01% |
| aprB | 0.00% | 0.00% | 0.00% | 0.00% | 0.01% | 0.01% | 0.02% | 0.02% | 0.00% | 0.00% | 0.00% | 0.00% | 0.00% | 0.00% | 0.00% | 0.00% | 0.00% | 0.00% | 0.00% | 0.00% | 0.00% | 0.00% | 0.00% | 0.00% | 0.00% | 0.00% | 0.00% | 0.00% | 0.01% | 0.00% |
| dsrA | 0.00% | 0.00% | 0.00% | 0.00% | 0.01% | 0.01% | 0.02% | 0.02% | 0.00% | 0.00% | 0.00% | 0.00% | 0.00% | 0.00% | 0.00% | 0.00% | 0.00% | 0.00% | 0.00% | 0.00% | 0.00% | 0.00% | 0.00% | 0.00% | 0.00% | 0.00% | 0.00% | 0.00% | 0.01% | 0.01% |
| dsrB | 0.00% | 0.00% | 0.00% | 0.00% | 0.00% | 0.01% | 0.01% | 0.01% | 0.00% | 0.00% | 0.00% | 0.00% | 0.00% | 0.00% | 0.00% | 0.00% | 0.00% | 0.00% | 0.00% | 0.00% | 0.00% | 0.00% | 0.00% | 0.00% | 0.00% | 0.00% | 0.00% | 0.00% | 0.01% | 0.01% |
| dsrC_tusE_dsvC | 0.00% | 0.01% | 0.01% | 0.02% | 0.02% | 0.05% | 0.06% | 0.08% | 0.00% | 0.00% | 0.00% | 0.00% | 0.00% | 0.00% | 0.00% | 0.00% | 0.00% | 0.00% | 0.00% | 0.00% | 0.00% | 0.00% | 0.00% | 0.00% | 0.00% | 0.03% | 0.00% | 0.00% | 0.04% | 0.04% |
| dsrE | 0.00% | 0.00% | 0.00% | 0.00% | 0.00% | 0.01% | 0.01% | 0.01% | 0.00% | 0.00% | 0.00% | 0.00% | 0.00% | 0.00% | 0.00% | 0.00% | 0.00% | 0.00% | 0.00% | 0.00% | 0.00% | 0.00% | 0.00% | 0.00% | 0.00% | 0.00% | 0.00% | 0.00% | 0.01% | 0.01% |
| dsrF | 0.00% | 0.00% | 0.00% | 0.00% | 0.00% | 0.01% | 0.01% | 0.01% | 0.00% | 0.00% | 0.00% | 0.00% | 0.00% | 0.00% | 0.00% | 0.00% | 0.00% | 0.00% | 0.00% | 0.00% | 0.00% | 0.00% | 0.00% | 0.00% | 0.00% | 0.00% | 0.00% | 0.00% | 0.01% | 0.01% |
| dsrH | 0.00% | 0.00% | 0.00% | 0.00% | 0.00% | 0.01% | 0.01% | 0.01% | 0.00% | 0.00% | 0.00% | 0.00% | 0.00% | 0.00% | 0.00% | 0.00% | 0.00% | 0.00% | 0.00% | 0.00% | 0.00% | 0.00% | 0.00% | 0.00% | 0.00% | 0.00% | 0.00% | 0.00% | 0.01% | 0.01% |
| fccB | 0.04% | 0.03% | 0.03% | 0.03% | 0.03% | 0.02% | 0.02% | 0.02% | 0.04% | 0.05% | 0.03% | 0.05% | 0.02% | 0.03% | 0.06% | 0.04% | 0.04% | 0.05% | 0.01% | 0.01% | 0.01% | 0.00% | 0.00% | 0.02% | 0.01% | 0.01% | 0.01% | 0.01% | 0.02% | 0.02% |
| sat_sulfate_adenylyltransferase | 0.00% | 0.00% | 0.00% | 0.00% | 0.01% | 0.01% | 0.01% | 0.02% | 0.00% | 0.00% | 0.00% | 0.00% | 0.00% | 0.00% | 0.00% | 0.00% | 0.00% | 0.00% | 0.00% | 0.00% | 0.00% | 0.00% | 0.00% | 0.00% | 0.00% | 0.00% | 0.00% | 0.00% | 0.01% | 0.01% |
| sor | 0.00% | 0.00% | 0.00% | 0.00% | 0.00% | 0.00% | 0.00% | 0.00% | 0.01% | 0.02% | 0.01% | 0.02% | 0.01% | 0.01% | 0.02% | 0.00% | 0.01% | 0.02% | 0.00% | 0.00% | 0.00% | 0.00% | 0.00% | 0.00% | 0.00% | 0.00% | 0.00% | 0.00% | 0.00% | 0.00% |
| SoxA | 0.02% | 0.02% | 0.01% | 0.02% | 0.02% | 0.02% | 0.02% | 0.03% | 0.02% | 0.02% | 0.01% | 0.02% | 0.01% | 0.01% | 0.02% | 0.02% | 0.02% | 0.02% | 0.01% | 0.02% | 0.01% | 0.01% | 0.01% | 0.01% | 0.01% | 0.02% | 0.01% | 0.02% | 0.03% | 0.02% |
| SoxB | 0.02% | 0.02% | 0.02% | 0.03% | 0.02% | 0.01% | 0.02% | 0.02% | 0.02% | 0.02% | 0.01% | 0.02% | 0.01% | 0.01% | 0.02% | 0.02% | 0.01% | 0.02% | 0.01% | 0.01% | 0.01% | 0.00% | 0.00% | 0.02% | 0.00% | 0.01% | 0.01% | 0.01% | 0.02% | 0.01% |
| SoxC | 0.02% | 0.02% | 0.02% | 0.01% | 0.01% | 0.00% | 0.00% | 0.00% | 0.02% | 0.02% | 0.01% | 0.02% | 0.01% | 0.01% | 0.02% | 0.02% | 0.02% | 0.02% | 0.01% | 0.01% | 0.00% | 0.00% | 0.00% | 0.01% | 0.00% | 0.01% | 0.01% | 0.01% | 0.01% | 0.01% |
| SoxX | 0.02% | 0.01% | 0.01% | 0.02% | 0.02% | 0.02% | 0.02% | 0.03% | 0.02% | 0.02% | 0.01% | 0.02% | 0.01% | 0.01% | 0.02% | 0.02% | 0.02% | 0.02% | 0.01% | 0.02% | 0.01% | 0.01% | 0.00% | 0.01% | 0.01% | 0.02% | 0.01% | 0.02% | 0.03% | 0.02% |
| SoxY | 0.03% | 0.03% | 0.03% | 0.03% | 0.02% | 0.02% | 0.02% | 0.03% | 0.02% | 0.02% | 0.01% | 0.02% | 0.01% | 0.01% | 0.02% | 0.02% | 0.02% | 0.02% | 0.01% | 0.01% | 0.01% | 0.00% | 0.00% | 0.02% | 0.00% | 0.01% | 0.01% | 0.01% | 0.02% | 0.01% |
| SoxZ | 0.04% | 0.04% | 0.03% | 0.04% | 0.03% | 0.01% | 0.02% | 0.02% | 0.02% | 0.02% | 0.01% | 0.02% | 0.01% | 0.01% | 0.02% | 0.02% | 0.02% | 0.02% | 0.01% | 0.01% | 0.01% | 0.00% | 0.00% | 0.02% | 0.00% | 0.01% | 0.01% | 0.01% | 0.02% | 0.02% |
| sulfur_dioxygenase_sdo | 0.02% | 0.02% | 0.02% | 0.02% | 0.02% | 0.04% | 0.04% | 0.04% | 0.04% | 0.04% | 0.05% | 0.05% | 0.05% | 0.05% | 0.04% | 0.04% | 0.04% | 0.04% | 0.04% | 0.06% | 0.07% | 0.02% | 0.03% | 0.01% | 0.05% | 0.05% | 0.05% | 0.05% | 0.05% | 0.04% |
| thiosulfate_reductase_phsA | 0.00% | 0.00% | 0.00% | 0.00% | 0.00% | 0.00% | 0.00% | 0.00% | 0.00% | 0.00% | 0.00% | 0.00% | 0.00% | 0.00% | 0.00% | 0.00% | 0.00% | 0.00% | 0.00% | 0.00% | 0.00% | 0.00% | 0.00% | 0.00% | 0.00% | 0.00% | 0.00% | 0.00% | 0.00% | 0.00% |

**Table S6.** Nitrogen speciation (N-$NO_3^-$, N-$NO_2^-$ and N-$NH_4^+$ (unfiltered; mmol/L)), total metal (Ni, Cu, Fe, Zn) ($0.45_{\mu m}$ and 0.2 $_{\mu m}$ (as indicated) **;** mg/L**)** and total dissolved organic carbon (DOC) ($0.45_{\mu m}$**,;** mmol/L) across time (2015-2018) of the tailings impoundment waters. All samples are shown as the average ± standard deviation with the exception of metal analysis where only averages are given.

| Date | Depth (m) | N-$NO_3^-$ mmol/L | N-$NO_2^-$ mmol/L | N-$NH_4^+$ mmol/L | DOC mmol/L |
|---|---|---|---|---|---|
| March 2015 | 0 | - | - | - | 0.45 ± 0.05 |
| | 0.75 | - | - | - | 0.28 ± 0.01 |
| | 21 | - | - | - | 0.34 ± 0.01 |
| May 2015 | 2.5 | - | - | - | 0.23 ± 0.01 |
| | 3.5 | - | - | - | 0.37 ± 0.01 |
| | 15 | - | - | - | 0.24 ± 0.01 |
| July 10 2015 | 2 | - | - | - | 0.29 ± 0.02 |
| | 4 | - | - | - | 0.26 ± 0.01 |
| | 30 | - | - | - | 0.23 ± 0.01 |
| July 22 2015 | 1 | - | - | - | 0.16 ± 0.01 |
| | 3.5 | - | - | - | 0.14 ± 0.01 |
| | 10 | - | - | - | 0.21 ± 0.01 |
| August 2015 | 1 | - | - | - | 0.52 ± 0.03 |
| | 2.8 | - | - | - | 0.15 ± 0.003 |
| | 26 | - | - | - | 0.26 ± 0.01 |
| Sept 2015 | 1 | - | - | 0.16 ± 0.006 | 0.10 ± 0.01 |
| | 7 | - | - | 0.16 ± 0.005 | 0.2 ± 0.001 |
| | 33 | - | - | 0.16 ± 0.008 | 0.2 ± 0.01 |
| Nov 2015 | 1.3 | - | - | 0.13 ± 0.0036 | 0.11 ± 0.003 |
| May 2016 | 1 | - | - | 0.12 ± 0.03 | 0.23 ± 0.05 |
| | 6 | - | - | 0.14 ± 0.003 | 0.24 ± 0.02 |
| July 2016 | 0.5 | - | - | 0.16 ± 0.003 | 0.2 ± 0.2 |
| | 1.5 | - | - | 0.16 ± 0.01 | 0.26 ± 0.4 |
| | 10 | - | - | 0.16 ± 0.012 | 0.2 ± 0.3 |
| Sept 2016 | 1 | - | - | 0.13 ± 0.003 | 0.22 ± 0.01 |
| | 6 | - | - | 0.13 ± 0.005 | 0.24 ± 0.01 |
| | 32 | - | - | 0.12 ± 0.003 | 0.24 ± 0.01 |
| May 2017 | 0.5 | 0.098 ± 0.004 | 0.003 ± 0.0002 | 0.06 ± 0.01 | 0.28 ± 0.02 |
| June 2017 | 0.5 | 0.114 ± 0.004 | 0.004 ± 0.0001 | 0.09 ± 0.006 | 0.19 ± 0.003 |
| | 2 | 0.105 ± 0.004 | 0.012 ± 0.0006 | 0.09 ± 0.009 | 0.3 ± 0.01 |
| | 10 | 0.115 ± 0.004 | 0.037 ± 0.0007 | 0.09 ± 0.006 | 0.32 ± 0.004 |
| July 2017 | 0.5 | 0.07 ± 0.004 | 0.02 ± 0.0004 | 0.16 ± 0.006 | 0.42 ± 0.01 |
| | 2 | 0.03 ± 0.007 | 0.031 ± 0.002 | 0.16 ± 0.009 | 0.42 ± 0.02 |
| | 10 | 0.068 ± 0.007 | 0.039 ± 0.0008 | 0.15 ± 0.02 | 0.59 ± 0.01 |
| August | 0.5 | 0.073 ± 0.005 | 0.017 ± 0.0001 | 0.14 ± 0.001 | 0.39 ± 0.01 |

| | | | | | |
|---|---|---|---|---|---|
| 2017 | 2 | 0.073 ± 0.004 | 0.017 ± 0.0006 | 0.15 ± 0.003 | 0.38 ± 0.01 |
| | 10 | 0.058 ± 0.004 | 0.023 ± 0.0002 | 0.13 ± 0.06 | 0.35 ± 0.01 |
| Sept 2017 | 0.5 | 0.042 ± 0.004 | 0.004 ± 0.0001 | 0.17 ± 0.006 | 1.32 ± 0.03 |
| | 2 | 0.039 ± 0.001 | 0.003 ± 0.0001 | 0.17 ± 0.003 | NA |
| | 10 | 0.019 ± 0.001 | 0.002 ± 0.0002 | 0.16 ± 0.003 | NA |
| Nov 2017 | 0.5 | 0.035 ± 0.004 | 0.005 ± 0.0002 | 0.16 ± 0.006 | NA |
| | 10 | 0.035 ± 0.011 | 0.005 ± 0.0005 | 0.16 ± 0.009 | NA |
| May 2018 | 0.5 | 0.068 ± 0.003 | 0.025 ± 0.0007 | 0.13 ± 0.006 | 1.01 ± 0.02 |
| | 2.8 | 0.105 ± 0.004 | 0.022 ± 0.002 | 0.13 ± 0.003 | 0.36 ± 0.01 |
| | 21 | 0.079 ± 0.017 | 0.038 ± 0.0001 | 0.15 ± 0.003 | 0.29 ± 0.01 |
| June 2018 | 0.5 | 0.102 ± 0.004 | 0.007 ± 0.0001 | 0.15 ± 0.005 | 0.57 ± 0.02 |
| July 11 2018 | 0.5 | 0.018 ± 0.001 | 0.018 ± 0.003 | 0.12 ± 0.1 | 0.71 ± 0.01 |
| | 2.8 | 0.017 ± 0.004 | 0.021 ± 0.001 | 0.15 ± 0.003 | 0.87 ± 0.02 |
| | 10 | 0.005 ± 0.004 | 0.026 ± 0.001 | 0.15 ± 0.006 | 0.31 ± 0.01 |
| July 23 2018 | 0.5 | 0.036 ± 0.004 | 0.011 ± 0.0001 | 0.17 ± 0.01 | 0.96 ± 0.02 |
| August 2018 | 0.5 | 0.05 ± 0.004 | 0.010 ± 0.003 | 0.16 ± 0.01 | 0.47 ± 0.01 |
| | 2.5 | 0.156 ± 0.056 | 0.015 ± 0.001 | 0.16 ± 0.01 | 0.34 ± 0.002 |
| | 10 | 0.018 ± 0.004 | 0.015 ± 0.001 | 0.16 ± 0.01 | 0.31 ± 0.01 |

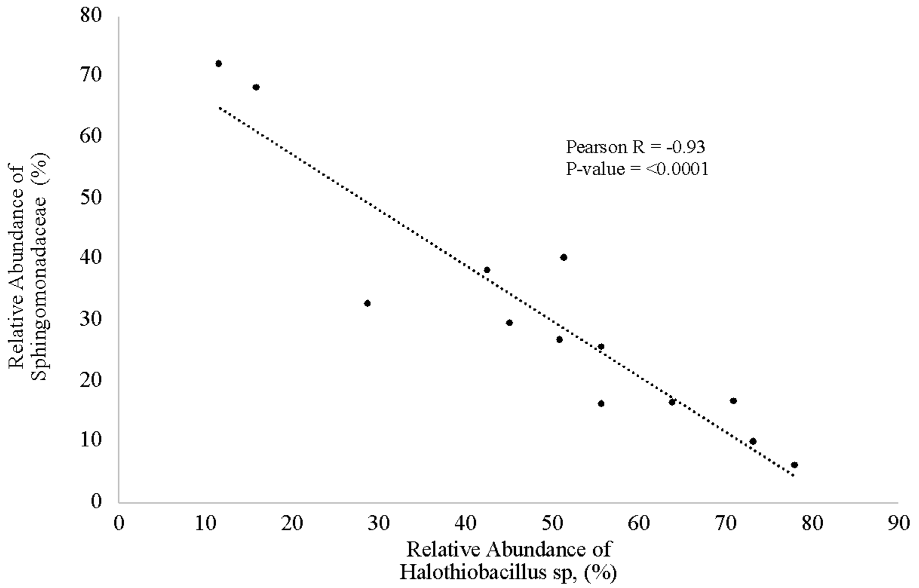


**Fig. S2 |** Correlational analysis between the relative abundances of *Halothiobacillus* sp. and the *Sphingomonadaceae* family in 2016/2017 waters.

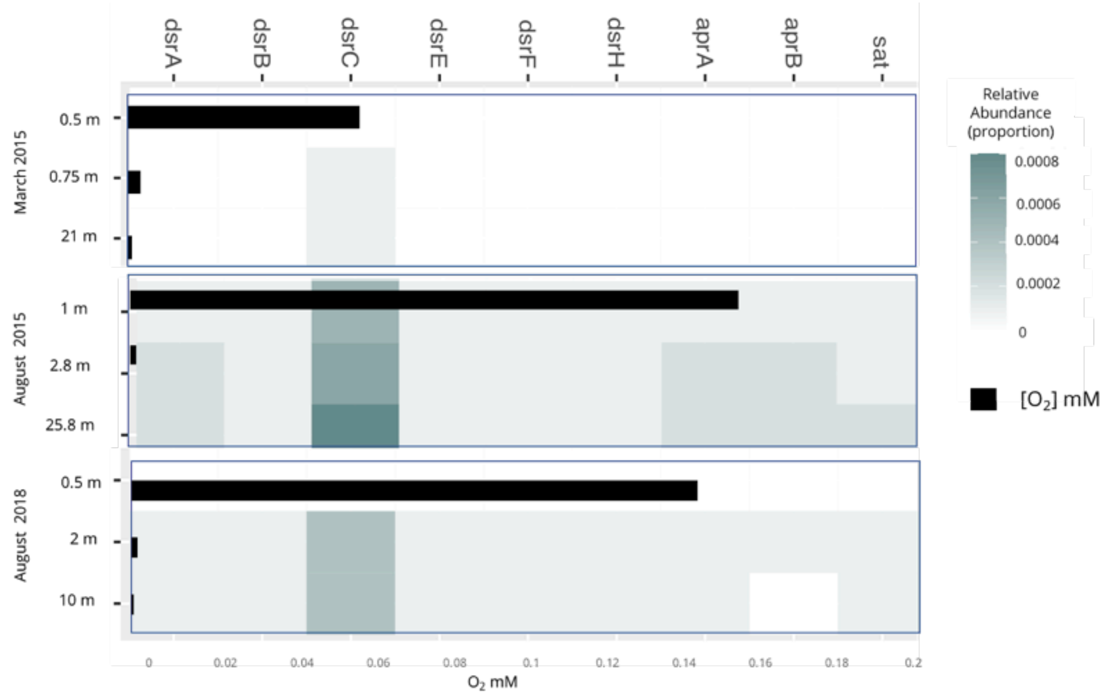


**Figure S3 |** Heat map of rDSR pathway gene relative abundances (dsrABCEFH, aprAB, sat) in March 2015, August 2015 and August 2018 through depth and overlaid oxygen concentrations (mM) (black bars) at corresponding depths.
